# Gene-teratogen interactions influence the penetrance of birth defects by altering Hedgehog signaling strength

**DOI:** 10.1101/2021.06.23.449683

**Authors:** Jennifer H. Kong, Cullen B. Young, Ganesh V. Pusapati, F Hernán Espinoza, Chandni B. Patel, Francis Beckert, Sebastian Ho, Bhaven B. Patel, George C. Gabriel, L. Aravind, J Fernando Bazan, Teresa M. Gunn, Cecilia W. Lo, Rajat Rohatgi

## Abstract

Birth defects result from interactions between genetic and environmental factors, but the mechanisms remain poorly understood. We find that mutations and teratogens interact in predictable ways to cause birth defects by changing target cell sensitivity to Hedgehog (Hh) ligands. These interactions converge on a membrane protein complex, the MMM complex, that promotes degradation of the Hh transducer Smoothened (SMO). Deficiency of the MMM component MOSMO results in elevated SMO and increased Hh signaling, causing multiple birth defects. *In utero* exposure to a teratogen that directly inhibits SMO reduces the penetrance and expressivity of birth defects in *Mosmo^-/-^* embryos. Additionally, tissues that develop normally in *Mosmo^-/-^* embryos are refractory to the teratogen. Thus, changes in the abundance of the protein target of a teratogen can change birth defect outcomes by quantitative shifts in Hh signaling. Consequently, small molecules that re-calibrate signaling strength could be harnessed to rescue structural birth defects.

## INTRODUCTION

Six percent of newborns suffer from structural birth defects, leading to 8 million cases a year worldwide (Christianson et al., 2005). Many of these structural defects require surgical intervention early in life and lead to adverse long-term health consequences. The underlying mechanisms driving birth defects remain unknown in a majority of cases. Complex interactions between genetic and environmental factors are thought to shift morphogen signaling beyond the threshold required for normal developmental patterning (Beames and Lipinski, 2020; Finnell, 1999; Krauss and Hong, 2016). However, in most cases the specific molecular mechanisms remain poorly understood. Penetrance and expressivity of birth defects, both between embryos and between tissues, remains unpredictable and confounds identification of causal factors. Improved understanding of molecular mechanisms is critical to developing strategies to alleviate the significant public health burden of birth defects.

The Hedgehog (Hh) pathway is one of a handful of signaling systems that regulate developmental patterning and morphogenesis of many tissues, including the face, limbs, heart, lungs, brain, and spinal cord (McMahon et al., 2003). Developing tissues are often exquisitely sensitive to the precise amplitude of Hh signaling. Even small changes in signaling strength can cause birth defects in mice and humans (Nieuwenhuis and Hui, 2005). Hh ligands are considered classical morphogens, secreted molecules that direct cell-fate choices in a dose-dependent manner (Lee et al., 2016). Temporal and spatial gradients of Hh ligands are translated into intracellular gradients of activity of the GLI transcription factors in target cells (Harfe et al., 2004; Jacob and Briscoe, 2003; Stamataki et al., 2005). Varying Hh signaling strength leads target cells to adopt different cell fates (Dessaud et al., 2008). Given the centrality of morphogen gradients in developmental patterning, considerable research effort has focused on understanding how they are established in tissues. However, this ligand-centric view of patterning is incomplete. Specific signaling mechanisms function in target cells to regulate their sensitivity to morphogens. Indeed, cell fate decisions often depend on both the extracellular concentration of ligands and the reception sensitivity of target cells to these ligands. A prominent example of such a mechanism can be found in the WNT pathway: cell-surface levels of Frizzled receptors (and consequently WNT sensitivity) is controlled by the transmembrane E3 ubiquitin ligases ZNRF3 and RNF43 (de Lau et al., 2014). These E3 ligases are themselves controlled by R-spondins, secreted ligands that play central roles in both pattern formation during development and in postnatal tissue homeostasis.

Our analysis of a gene called *Mosmo* (<u>Mo</u>dulator of <u>Smo</u>othened) led us to uncover a cell-surface pathway that regulates the sensitivity to Hh ligands and consequently the development of multiple tissues. Mouse genetic analysis revealed that *Mosmo* uniquely functions to tune the Hh signaling gradient in target cells by promoting the degradation of Smoothened (SMO), a 7-pass transmembrane protein that carries the Hh signal across the plasma membrane. Interestingly, mutations in *Mosmo* (which increase SMO protein abundance) influence penetrance, expressivity and tissue specificity of birth defects caused by an exogenous teratogen that directly inhibits SMO activity. These findings show that the penetrance of birth defects can be modulated by gene-environment interactions that alter ligand sensitivity in the Hh pathway.

## RESULTS

### MOSMO is required for embryonic development

*Mosmo* is a novel gene we initially identified in a loss-of-function CRISPR screen conducted in NIH/3T3 fibroblasts designed to find negative regulators of Hedgehog (Hh) signaling (Pusapati et al., 2018). *Mosmo* encodes an 18.2 kDa 4-pass transmembrane protein of unknown function. Depletion of MOSMO in cultured cells results in increased accumulation of the Hh transducer Smoothened (SMO) on the plasma membrane and the primary cilium membrane, resulting in hyper-responsiveness to Hh ligands. *Mosmo* is widely expressed in mouse embryos, based on *in situ* hybridization (**Figure S1A**) and the analysis of published single-cell RNAseq data (Pijuan-Sala et al., 2019) (**Figure S1B**). To understand the developmental roles of MOSMO, we used CRISPR/Cas9 genome editing to generate mice carrying null alleles of *Mosmo* (**Figure S2A and S2B**). While *Mosmo*^+/-^ mice developed normally, no live *Mosmo*^-/-^ pups were recovered from heterozygous intercrosses (**Figure 1A** **and Table S1**). Most *Mosmo*^-/-^ embryos die by gestational day 14.5 (e14.5) (**Figure 1A** **and Table S1**). We conclude that the function of MOSMO is essential for embryonic development.

**Figure 1:**
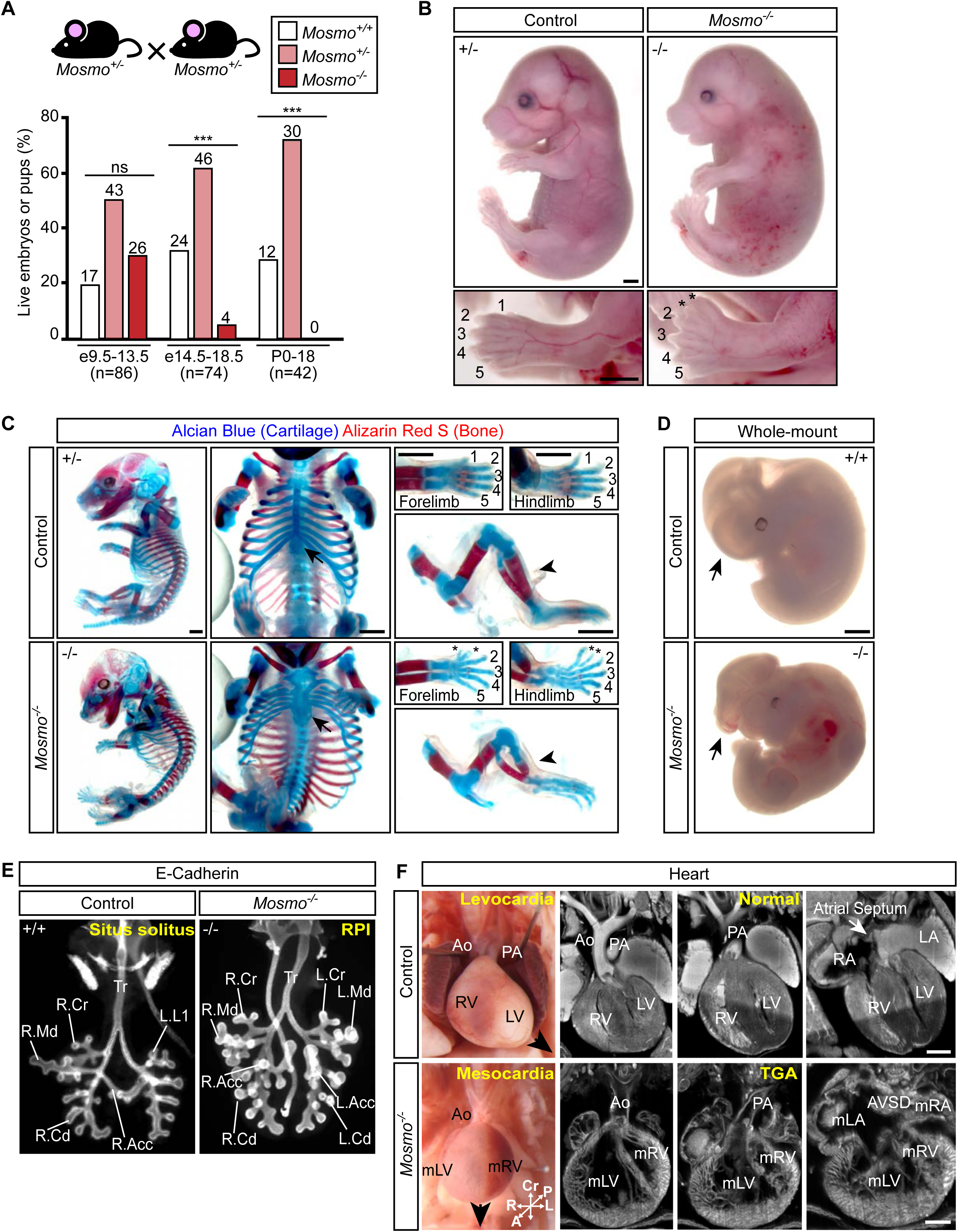
Loss of *Mosmo* results in embryonic lethality and developmental defects. **(A)** Viability of offspring derived from *Mosmo^+/-^* x *Mosmo^+/-^* crosses at the indicated developmental stages. Statistical significance was determined by the chi-square test; not-significant (ns) > 0.05 and \*\*\**p*-value ≤ 0.001; *n* = number of live embryos collected. See Table S1 for full details. **(B)** Whole-mount images of e16.5 control (*Mosmo^+/-^*) and *Mosmo^-/-^* littermates show that the latter have edema and preaxial polydactyly. See Table S2 for a detailed list of phenotypes in each embryo analyzed. Scale bars are 1 mm. **(C)** Skeletons from e16.5 control (*Mosmo^+/-^*) and *Mosmo^-/-^* littermates stained with Alcian Blue and Alizarin Red S to visualize cartilage and calcified bone, respectively. Polydactyly (asterisks), sternal clefting (middle column, arrows) and tibial truncation (tibial hemimelia, right column, arrowheads) were observed in *Mosmo^-/-^* embryos. Scale bars are 1 mm. **(D)** Whole-mount images of e11.5 control (*Mosmo^+/+^*) and *Mosmo^-/-^* littermates show that the latter suffer from exencephaly (arrow). Scale bar is 1 mm. **(E)** Whole-mount lungs (ventral view) of e12.5 control (*Mosmo^+/+^*) and *Mosmo*^-/-^ embryos immunostained for E-cadherin to show the airway epithelium and allow for detailed branching analysis. Normal mouse lungs have one lobe on the left (L.L1) and four lobes on the right (R.Acc, right accessory; R.Cd, right caudal; R.Cr, right cranial; R.Md, right middle). *Mosmo*^- /-^ lungs exhibit right pulmonary isomerism (RPI), a duplication of the right lung morphology on the left side (L.Acc, left accessory; L.Cd, left caudal; L.Cr, left cranial; L.Md, left middle). Further details are provided in Table S3. **(F)** *Mosmo^-/-^* mutant embryos exhibit complex CHDs associated with abnormal left-right patterning of the heart. Necropsy and Episcopic Confocal Fluorescence Microscopy (ECM) images of representative embryonic hearts from e16.5 *Mosmo^+/+^* (control, top) and *Mosmo*^-/-^ embryos (bottom). The *Mosmo*^-/-^ heart has no apparent apex (down arrow), indicating mesocardia, with the aorta (Ao) abnormally positioned anteriorly. The Ao is situated anterior to the pulmonary artery (PA) and inserted into the morphological right ventricle (mRV) situated on the body’s left, while the pulmonary artery (PA) emerges from the morphological left ventricle (mLV) positioned on the body’s right, findings diagnostic of transposition of the great arteries (TGA). Other findings include non-compaction of the ventricular myocardium and an unbalanced atrioventricular septal defect (AVSD) with symmetric insertions of both inferior and superior vena cava suggesting right atrial isomerism (mRA). Scale bar is 0.5 mm. A detailed list of cardiac phenotypes observed in each embryo can be found in Table S2.

### *Mosmo* is required for proper left-right patterning and heart, limb and lung development

*Mosmo* deficiency results in developmental defects across many organ systems. *Mosmo*^-/-^ embryos have preaxial polydactyly in both forelimbs and hindlimbs (**Figures 1B and 1C)**. Whole-mount skeletal staining revealed additional skeletal defects, including a split sternum (**Figure 1C, arrows**) and truncated tibia (**Figure 1C, arrowheads**). A subset of *Mosmo*^-/-^ embryos exhibited exencephaly (**Figure 1D**). Detailed necropsy examination of the internal anatomy revealed that all *Mosmo*^-/-^ embryos had heterotaxy, discordant patterning of the left-right body axis manifested as abnormalities in lung lobation and abnormal left-right positioning of multiple visceral organs including the heart, stomach, spleen, and pancreas (**Table S2**). Analysis of the early lung branching pattern indicated that most *Mosmo*^-/-^ embryos have either complete or partial right pulmonary isomerism, a duplication of the right lung morphology on the left side (**Figures 1E** **and S3A, Table S3**). Analysis of the developing heart using episcopic confocal fluorescence microscopy (ECM) revealed that all *Mosmo*^-/-^ embryos have complex congenital heart defects (CHDs). The most common CHDs observed are transposition of the great arteries (TGA) and atrioventricular septal defects (AVSDs) (**Figure 1F** **and Table S2**). TGA and AVSDs are classified as “critical” heart defects in human patients since they require surgical intervention soon after birth. *Mosmo*^-/-^ embryos likely die *in utero* due to these complex structural heart defects.

### *Mosmo^-/-^* phenotypes are correlated with elevated Hh signaling activity

To understand the etiology of the birth defects observed in *Mosmo*^-/-^ embryos, we focused on the Hh signaling pathway because *Mosmo* was originally identified as an attenuator of Hh signaling in our CRISPR screens (Pusapati et al., 2018), and many of the *Mosmo*^-/-^ phenotypes (i.e. polydactyly and exencephaly) can be caused by elevated Hh signaling (Hui and Joyner, 1993). To assess Hh signaling activity in *Mosmo^-/-^* cells, primary mouse embryonic fibroblasts (pMEFs) were isolated and treated with varying concentrations of Sonic Hedgehog (SHH), a secreted ligand that initiates Hh signaling in target cells. Compared to cells from wild-type littermate controls, Hh signaling was elevated in *Mosmo^-/-^* pMEFs (**Figure 2A**). A low concentration of SHH (1 nM) that failed to fully activate expression of the Hh target gene *Gli1* in wild-type pMEFs was sufficient to maximally activate *Gli1* in *Mosmo*^-/-^ pMEFs.

**Figure 2:**
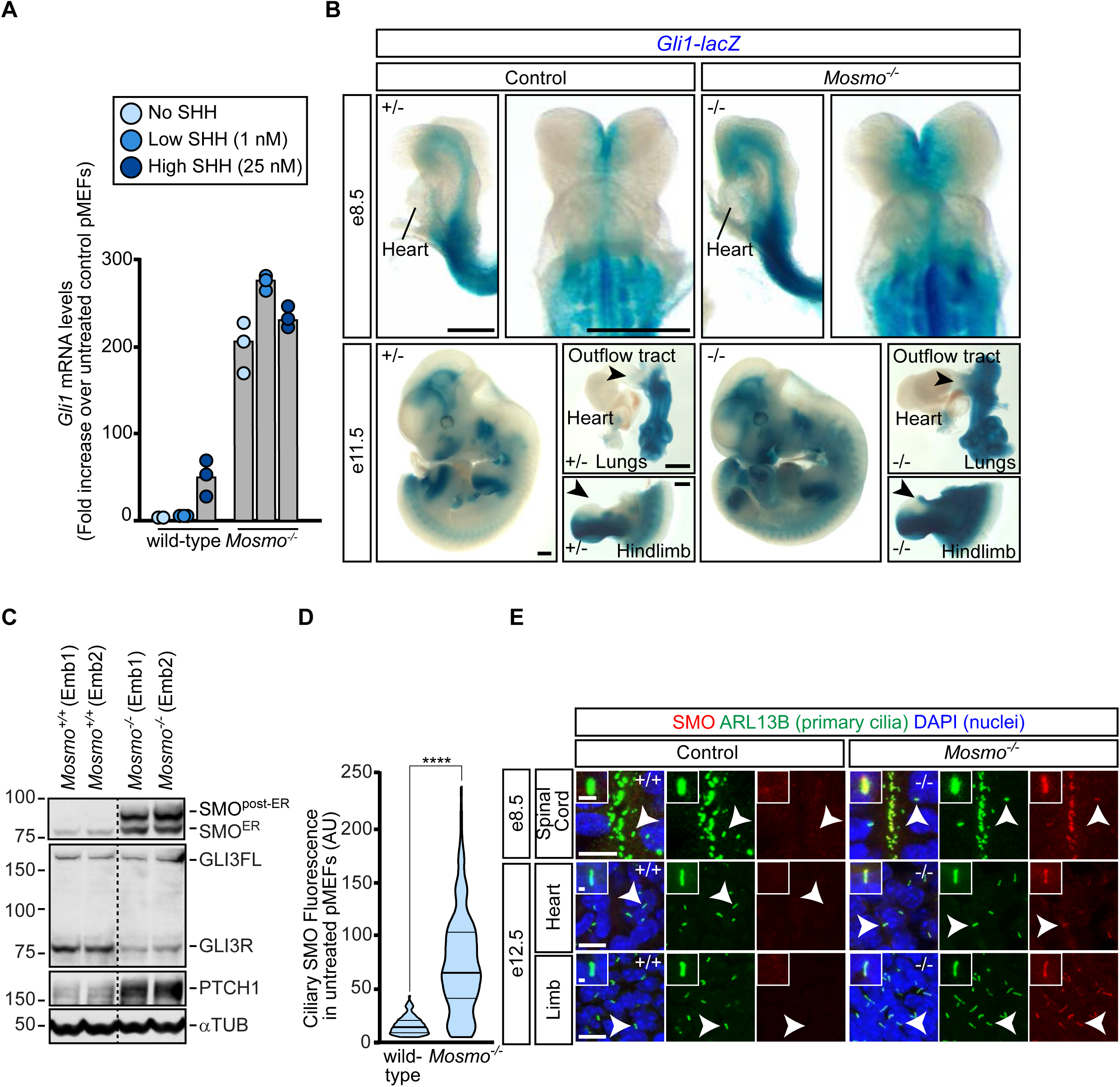
Loss of *Mosmo* results in elevated Hh signaling activity. Hh signaling activity in primary mouse embryonic fibroblasts (pMEFs) and embryonic tissues was assessed using expression of *Gli1*, a direct Hh target gene, or accumulation of SMO in primary cilia. **(A)** *Gli1* mRNA abundance in wild-type (*Mosmo^+/+^*) and *Mosmo^-/-^* pMEFs was measured using qRT-PCR. Bars denote the median *Gli1* mRNA values derived from the two to three individual measurements shown as circles. **(B)** X-gal staining was used to visualize *Gli1-lacZ* expression in whole-mount preparations of mouse embryos. Arrowheads denote areas of elevated Hh signaling activity in the cardiac outflow tract and anterior hindlimb. Scale bars are 0.5 mm. **(C)** Immunoblot used to measure protein abundance of SMO, GLI3, PTCH1, and ɑTUB (a loading control) in whole embryo lysates prepared from e12.5 wild-type (*Mosmo^+/+^*) and *Mosmo^-/-^* littermates. **(D and E)** Immunofluorescence (IF) was used to measure SMO abundance (red) in primary cilia (ARL13B, green) in pMEFs (D) and various embryonic tissues (E). Violin plots in (D) summarize SMO fluorescence data from ∼100-150 cilia, with horizontal lines denoting the median and interquartile range. Statistical significance was determined by the Mann-Whitney test; \*\*\*\**p*-value ≤ 0.0001. (E) DAPI (blue) marks nuclei. White arrows denote the primary cilium enlarged in the inset. Scale bars, 10 µm in merged panels and 1 µm in zoomed displays.

To assess levels of Hh signaling activity in *Mosmo^-/-^* embryos, we crossed *Mosmo^+/-^* mice with *Gli1^lacZ/+^* mice, an extensively used Hh reporter line in which a lacZ transgene (encoding β-galactosidase) was inserted into the first coding exon of *Gli1* (Bai et al., 2002). The *Gli1-lacZ* expression pattern recapitulated endogenous *Gli1* expression and the expression of other Hh target genes like *Ptch1* (Goodrich et al., 1997; Guzzetta et al., 2020; Hui et al., 1994). At e8.5 and e9.5, *Gli1-lacZ* was expressed in the neural tube, somites, and secondary heart field (SHF) (**Figures 2B** **and S3B**) (Guzzetta et al., 2020). At e11.5, *Gli1* reporter activity was observed in the brain (telencephalon and diencephalon), spinal cord, limbs, lungs, frontonasal processes, and pharyngeal endoderm (**Figures 2B** **and S3B**). Compared to littermate controls (*Mosmo^+/+^* and *Mosmo^+/-^*), *Gli1-lacZ* expression was elevated in *Mosmo^-/-^* embryos at all embryonic ages analyzed. The expansion of *Gli1* expression is notable in the developing limb and cardiac outflow tract (**Figure 2B** **and S3C**), consistent with the finding of polydactyly and outflow tract related heart defects in *Mosmo^-/-^* embryos (**Table S2**) (Goddeeris et al., 2007). Consistent with the elevation of *Gli1* expression, we observed an increase in PTCH1 (encoded by a direct Hh target gene) and a decrease GLI3R (the major Hh transcriptional repressor) in *Mosmo^-/-^* whole embryo lysates, showing that *Mosmo* deficiency results in increased Hh signaling activity *in vivo* (**Figure 2C**).

Hh signal transmission across the plasma membrane requires the SHH-induced accumulation of SMO in the membrane of the primary cilium (Corbit et al., 2005; Rohatgi et al., 2007). The loss of *Mosmo* led to the constitutive, high-level accumulation of SMO in the ciliary membrane in pMEFs (**Figure 2D**) and all embryonic tissues analyzed (**Figure 2E**). We also observed an increase in SMO protein abundance in *Mosmo^-/-^* embryo lysates (**Figure 2C**). This increase was dramatic for the SMO protein band that migrated more slowly in the SDS-PAGE gel, which represents the population of SMO that has transversed the endoplasmic reticulum (post-ER) and acquired glycan modifications attached in the Golgi. We conclude that *Mosmo* functions to attenuate Hh signaling strength in both cells and embryos by reducing SMO levels in the ciliary membrane. Re-expression of MOSMO into *Mosmo^-/-^* cells restored both wild-type Hh signaling and ciliary SMO levels (**Figures S4A and S4B**).

### MOSMO interacts with MEGF8 and MGRN1 to form the MMM complex

The cellular phenotypes (elevated ciliary SMO and sensitivity to SHH) and developmental defects (polydactyly, heterotaxy and CHDs) seen in *Mosmo^-/-^* embryos were reminiscent of those caused by the loss of either *Megf8* or *Mgrn1* and *Rnf157*, components of a membrane-tethered E3 ubiquitin ligase complex that ubiquitinates SMO and accelerates its endocytosis and degradation (Kong et al., 2020) (**Figures 3A** **and S4C, Table S2**). To determine if MEGF8 and MOSMO are part of the same pathway, we compared Hh signaling activity and SMO abundance in cells lacking each gene individually (*Mosmo^-/-^* and *Megf8^-/-^* single knockouts) to cells lacking both (*Mosmo*^-/-^;*Megf8*^-/-^ double knockouts). The increase in cell-surface SMO and SHH sensitivity seen in *Mosmo*^-/-^;*Megf8*^-/-^ double knockout cells was comparable to that seen in *Mosmo^-/-^* and *Megf8*^-/-^ single knockout cells (**Figure S4D**). Similarly, the birth defects observed in *Mosmo^-/-^*;*Megf8^m/m^* double mutant mouse embryos were comparable in both penetrance and expressivity to *Mosmo^-/-^* and *Megf8^m/m^* single mutant embryos (**Figure 3A** **and Table S2**). Taken together, analysis in both cells and embryos suggested that *Mosmo* and *Megf8* belong to the same epistasis group and thus the proteins encoded by these genes likely belong to the same pathway.

**Figure 3:**
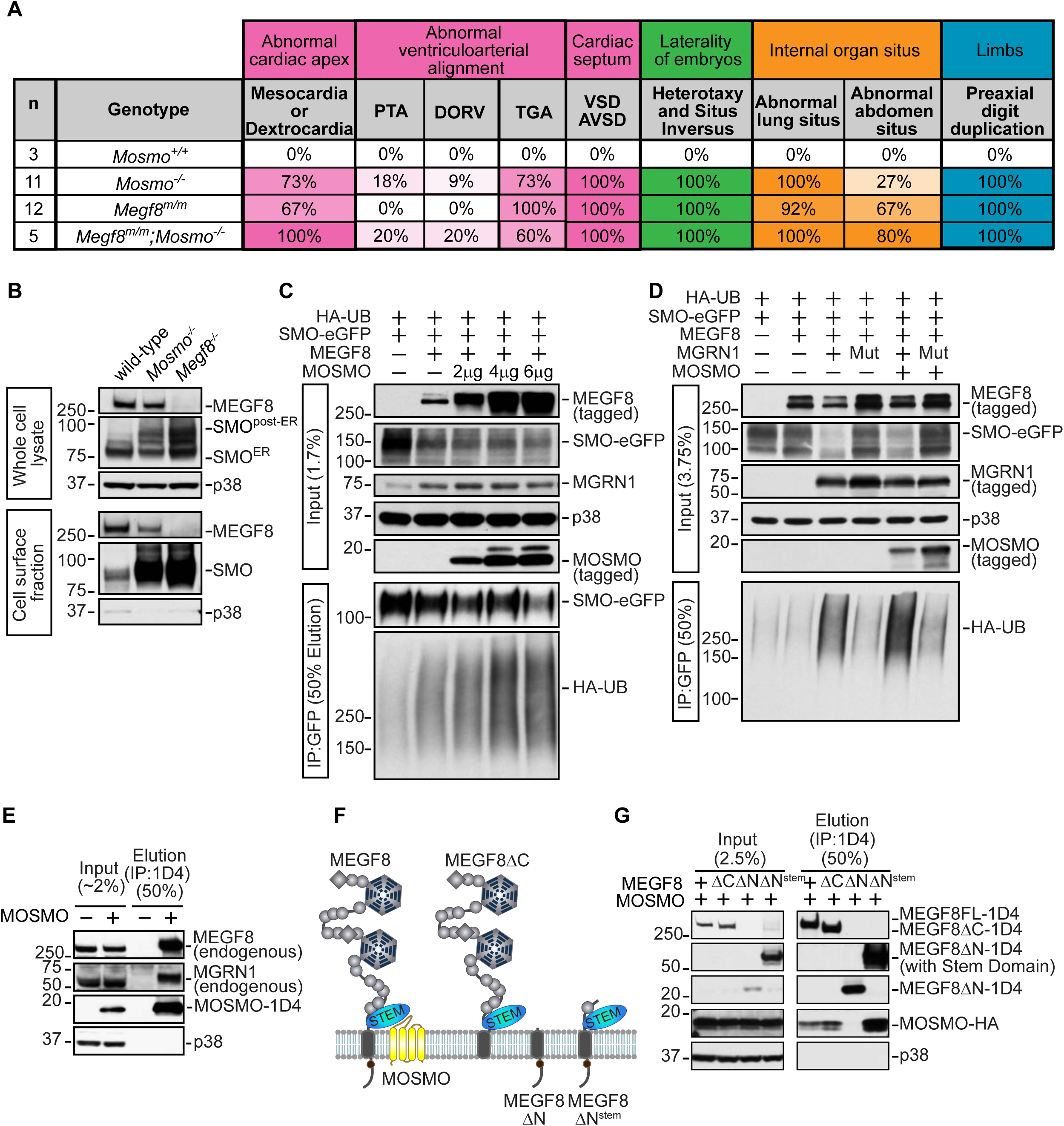
MOSMO interacts with MEGF8 and regulates its abundance at the cell surface. **(A)** Tabulation of phenotypes observed in e13.5-14.5 mouse embryos of the various genotypes as determined by necropsy and ECM imaging. AVSD, atrioventricular septal defect; DORV, double outlet right ventricle; PTA, persistent truncus arteriosus; TGA, transposition of the great arteries; VSD, ventricular septal defect. A detailed list of phenotypes observed in each embryo can be found in Table S2. The *Megf8^-/-^* phenotyping data is from our previous study (Kong et al., 2020). **(B)** Abundance of SMO and MEGF8 protein in whole cell lysates (top, 6.25% input) and on the cell-surface (bottom, cell surface biotinylation and streptavidin immunoprecipitation (IP), 50% elution) in NIH/3T3 cells of the indicated genotypes. The cytoplasmic protein p38 serves as a loading control. **(C-D)** SMO ubiquitination was assessed after transient co-expression of the indicated proteins in HEK293T cells, as described in our previous study (Kong et al., 2020). Cells were lysed under denaturing conditions, SMO was purified by IP, and the amount of HA-tagged Ubiquitin (HA-UB) covalently conjugated to SMO was assessed using immunoblotting with an anti-HA antibody. In (C), assays were done in the presence of endogenous MGRN1 and increasing amounts of transfected MOSMO. Assays in (D) co-transfected either wild-type MGRN1 or catalytically inactive MGRN1 (MGRN1^Mut^) to show that SMO ubiquitination was dependent on the function of MGRN1. **(E)** Endogenous MEGF8 and MGRN1 co-purified with 1D4-tagged MOSMO immunoprecipitated from *Mosmo^-/-^* NIH/3T3 cells stably expressing MOSMO-1D4 (**Figure S4A and S4B**). (**F**) Domain graphics depict the simplified modular architecture of wild-type MEGF8 (leftmost image) and its engineered variants (right three images). Grey beads denote the linearly-connected EGFL and PSI repeats, with interspersed 6-bladed β-propeller domains (hexagons) and CUB domains (diamonds), while the juxtamembrane Stem domain is a blue oval. The larger globular structures of the β-propeller, CUB and Stem folds all have closely spaced N- and C-termini, so they appear as pendant-like inserts into the long EGFL and PSI chain. A proposed mode of interaction between the Stem domain of MEGF8 and the extracellular domain of MOSMO is depicted (leftmost), based on modelling described in **Figure S5**. (**G**) A series of truncation mutants of MEGF8 (shown in **F**) were used to identify the MEGF8 domain that binds to MOSMO. The interaction between HA-tagged MOSMO and these 1D4-tagged MEGF8 variants was assessed by transient expression in HEK293T cells followed by an anti-1D4 IP and immunoblotting to measure the amount of co-precipitated MOSMO-HA (right).

A clue to the biochemical function of MOSMO came from the observation that the abundance of cell-surface MEGF8 was reduced in *Mosmo^-/-^* cells. Cell-surface biotinylation analysis demonstrated that the loss of MOSMO reduced MEGF8 (and concomitantly increased SMO) at the plasma membrane (**Figure 3B**). These results are consistent with the model that MOSMO promotes ubiquitination of SMO via the MEGF8-MGRN1 complex by increasing MEGF8 levels at the cell surface. Indeed, co-expression of MOSMO increased the ubiquitination of SMO by the MEGF8-MGRN1 complex in an assay reconstituted in HEK293T cells (**Figures 3C and 3D**). The influence of MOSMO on MEGF8 activity and localization led us to test the possibility of a physical interaction between the two proteins. Epitope-tagged MOSMO stably expressed in *Mosmo^-/-^* cells (**Figures S4A and S4B**) was co-immunoprecipitated with both endogenous MEGF8 and MGRN1 (**Figure 3E**).

The interaction between MEGF8 and MOSMO mapped to a previously unrecognized and unclassified juxtamembrane or “Stem” domain in MEGF8, so named because it is positioned at the extracellular end of the MEGF8 transmembrane helix and supports the rest of the large extracellular domain (**Figure 3F and 3G**). Using sensitive deep-learning-based structure prediction methods (Yang et al., 2020), the Stem domain is predicted to adopt a novel β-jellyroll fold topologically related to CUB and GOLD domains (**Figure S5A and S5B**) (Schaeffer et al., 2017). We propose that the Stem domain of MEGF8 docks to the compact extracellular β-sheet surface of MOSMO. Since MOSMO is distantly related to the claudins (Pusapati et al., 2018), we modelled this interaction based on the binding of clostridium perfringens enterotoxins (that fold as CUB-fold domains) to Claudin-3, -4, and -19 (Suzuki et al., 2017) (**Figure S5C**, left). Taken together, we propose that MOSMO, MEGF8 and MGRN1 together form a membrane-tethered E3 ligase complex (hereafter the “MMM complex”, **Figure S5C**, right) that regulates the strength of Hh signaling by regulating levels of SMO at the cell surface and primary cilium.

### *Mosmo^-/-^* limb phenotypes can be suppressed by the small-molecule SMO inhibitor vismodegib

As a component of the MMM complex, MOSMO attenuates Hh signaling activity in the developing embryo (**Figure 2A-C**) by clearing SMO from the cell surface and primary cilium (**Figure 2D and 2E**). However, MOSMO may also regulate other cellular pathways and processes. Thus, we sought to investigate whether the developmental defects (i.e., polydactyly and CHDs) observed in *Mosmo^-/-^* embryos (**Figure 1** **and** **Figure 3A**) were caused by elevated Hh signaling. We took the unconventional approach of using an FDA-approved small-molecule SMO antagonist (vismodegib) to reduce Hh signaling strength at critical periods of embryonic development. There are many advantages to using small-molecule inhibitors in an embryonic system. First, previous studies have shown that the SMO inhibitors cyclopamine and vismodegib are potent placenta-permeable teratogens that can induce embryonic defects in Hh-dependent tissues when delivered orally to the pregnant mother (Binns et al., 1963; Lipinski et al., 2008). Second, a small-molecule strategy allows us to selectively reduce Hh signaling during defined developmental periods to target events like limb digitation and heart looping. Lastly, by changing the treatment dose and frequency, a small-molecule inhibitor allows us to experimentally adjust Hh signaling activity as needed based on the phenotypic outcomes (Heyne et al., 2015). In a proof-of-concept experiment, we found that treatment of pregnant mice with vismodegib for about two days (e9.75-e11.5) reduced *Gli1-lacZ* expression in wild-type embryos when compared to untreated controls (**Figure 4A**).

**Figure 4:**
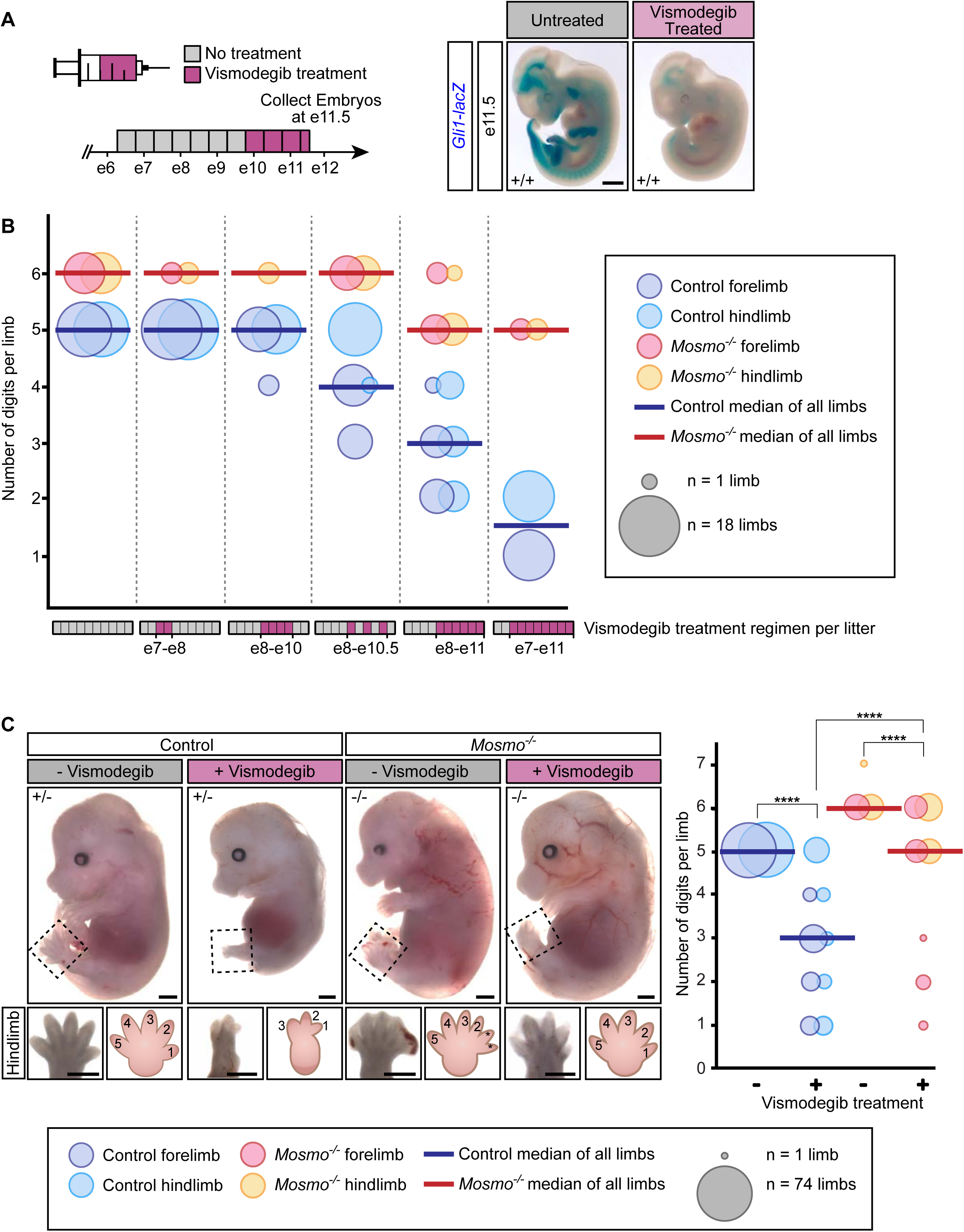
Reducing Hh signaling strength with the SMO rescues limb defects in *Mosmo^-/-^* embryos. **(A)** Vismodegib administered every 12 hours for ∼2 days from e9.75-e11.5 (diagrammed on left, with purple used to shade developmental windows for drug exposure) reduced *Gli1-lacZ* expression in wild-type embryos (right). Scale bar is 1 mm. **(B)** Numbers of digits per limb of e14.5 embryos from six individual litters exposed to increasing amounts vismodegib (represented by purple shading in the developmental time courses depicted on the x-axis). The area of each bubble is proportional to the number of limbs included in the analysis. Bold horizontal lines represent the median number of digits per limb in control (*Mosmo^+/+^* and *Mosmo^+/-^,* blue) and *Mosmo^-/-^* (red) embryos. **(C)** Representative images (left) of e14.5 control and *Mosmo^-/-^* embryos treated with or without vismodegib every 12 hours for three (e8.25- e11.25) or four (e7.25-e11.25) days. Scale bars are 1 mm. Bubble plot (right) used to depict the numbers of digits per limb of control (*Mosmo^+/+^* and *Mosmo^+/-^,* blue) and *Mosmo^-/-^* (red) embryos treated with vismodegib. Statistical significance was determined by the Kruskal-Wallis test; ****p value ≤ 0.0001.

*Shh* is transiently expressed along the posterior margin from ∼e9.5-e12 in the murine forelimb and ∼e10-e12.5 in the hindlimb (Büscher et al., 1997; Zhu et al., 2008). Hh signaling plays an established role in the anterior-posterior patterning of digits. Oligodactyly (digit loss) can be caused by exposure to a Hh antagonist or loss of *Shh* expression during a critical period of limb development (Heyne et al., 2015; Zhu et al., 2008). Conversely, preaxial polydactyly can arise from elevated Hh signaling activity caused by an increase in *Shh* or reduction in *Gli3* expression (Hill and Lettice, 2013; Hui and Joyner, 1993). To determine if the fully penetrant preaxial polydactyly observed in the *Mosmo^-/-^* embryos was due to elevated Hh signaling activity, pregnant dams from *Mosmo^+/-^* x *Mosmo^+/-^* crosses were exposed to vismodegib for varying durations of time and e14.5 embryos were collected to examine digit patterning. There were no defects in limb patterning in embryos that received no treatment or embryos that were treated with vismodegib before e9.5 (prior to *Shh* limb expression). However, when embryos were treated with vismodegib after e9.5, increasing the duration of drug treatment resulted in progressively greater oligodactyly (**Figure 4B**). The exquisitely graded nature of Hh signaling was vividly demonstrated by the striking dose-response relationship between vismodegib exposure and digit number. Interestingly, vismodegib treatment had a profoundly different impact on limb patterning even between embryos in the same litter (**Figure 4B**). While vismodegib caused severe oligodactyly in control (*Mosmo^+/+^* and *Mosmo^+/-^*) embryos, the same dose often corrected the polydactyly in *Mosmo^-/-^* embryos, resulting in embryos with normal limbs bearing five digits (**Figure 4B and 4C**). We conclude that the polydactyly observed in *Mosmo^-/-^* embryos is indeed due to an elevation of SMO activity since it can be reversed by a direct SMO antagonist (**Figure S6A**).

### *Mosmo^-/-^* cardiac phenotypes can be partially suppressed by SMO inhibitors

The ability of vismodegib to rescue the polydactyly phenotype led us to test its effects on the complex CHD phenotypes contributing to the lethality of *Mosmo^-/-^* embryos (**Figures 1A and 1F**). Conotruncal heart defects are a group of malformations that arise due to defects in outflow tract development. All of the *Mosmo^-/-^* embryos had conotruncal heart defects, most commonly Transposition of the Great Arteries (TGA) (**Figure 3A** **and Table S2**). Hh signaling plays a critical role in multiple aspects of outflow tract development including: (1) maintenance of cardiac progenitor proliferation and identity within the secondary heart field (which contributes to the outflow tract) (Dyer and Kirby, 2009; Rowton et al., 2020), (2) survival of migratory cardiac neural crest cells (Washington Smoak et al., 2005), and (3) proper septation of the outflow tract (Goddeeris et al., 2007). To determine if the conotruncal heart defects observed in the *Mosmo^-/-^* embryos are due to elevated Hh signaling, we first needed to identify the critical time window during gestation when outflow tract development is sensitive to vismodegib. Loss-of-function mutations in *Shh* result in a failure of the primitive truncus to properly divide into the aorta and pulmonary artery (Persistent Truncus Arteriosus, PTA) (Washington Smoak et al., 2005). Building on this information, we found that vismodegib administered from e7.25/e8.25 to e11.25 caused PTA in all control (*Mosmo^+/+^* and *Mosmo^+/-^*) embryos (**Figures 5A and 5B**). In contrast, *Mosmo^-/-^* embryos exposed *in utero* to the same vismodegib regimen developed did not develop PTA, suggesting that these mutant embryos were resistant to the SMO antagonist because of elevated SMO abundance (**Figure 5B**).

**Figure 5:**
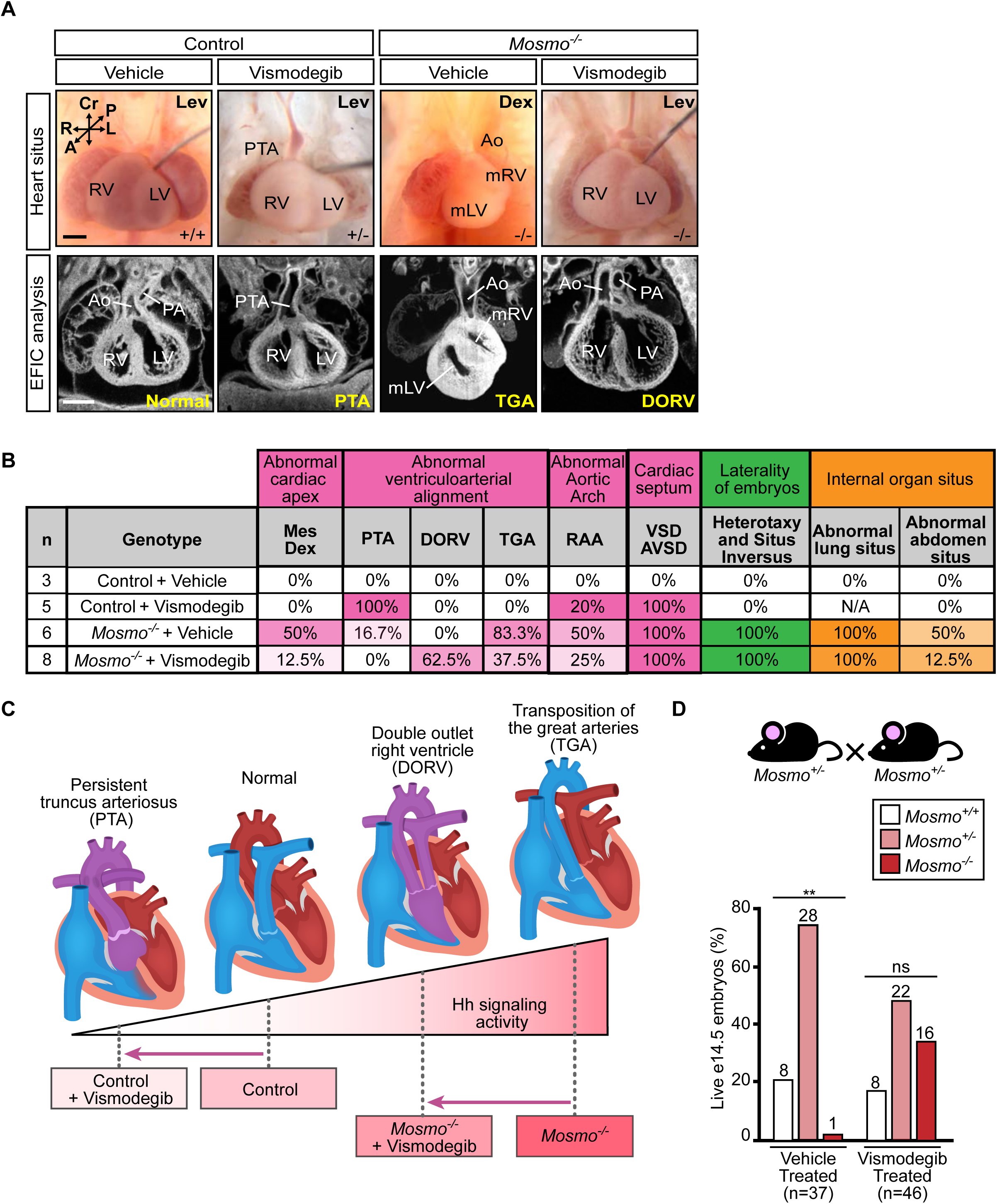
Reducing HH signaling strength partially rescues heart defects in *Mosmo^-/-^* embryos. **(A)** Representative necropsy (top row) and ECM (bottom row) images of embryonic hearts from e14.5 control (*Mosmo^+/+^* and *Mosmo^+/-^*) and *Mosmo^-/-^* embryos treated with vismodegib or a vehicle control. Vismodegib was administered every 12 hours for three (e8.25-e11.25) or four (e7.25-e11.25) days. Scale bar is 200 µm. **(B)** Summary of heart malformations and left-right patterning defects in e14.5 control (*Mosmo^+/+^* and *Mosmo^+/-^*) and *Mosmo^-/-^* mouse embryos treated with either vehicle or vismodegib every 12 hours for three (e8.25-e11.25) or four (e7.25-e11.25) days. Lung situs could not be determined (N/A) in vismodegib treated control embryos due to cystic and hypoplastic lungs. Ao, aorta; DORV, double outlet right ventricle; LV, left ventricle; mLV, morphological left ventricle; mRV, morphological right ventricle; PA, pulmonary artery; PTA, persistent truncus arteriosus; RV, right ventricle; TGA, transposition of the great arteries. A detailed list of phenotypes observed in each embryo can be found in Table S4. **(C)** A proposed model for how a combination of genotype and SMO inhibition by vismodegib influences Hh signaling strength and consequently ventriculoarterial alignment in developing embryos. **(D)** Viability of e14.5 embryos from *Mosmo^+/-^* x *Mosmo^+/-^* crosses treated with vehicle or vismodegib every 12 hours for three (e8.25-e11.25) or four (e7.25-e11.25) days. Statistical significance was determined by the chi-square test; not-significant (ns) > 0.05 and \*\**p*-value ≤ 0.01; *n* = number of live embryos collected. See Table S5 for full details.

Indeed, vismodegib administration actually improved the CHD phenotypes characteristically seen in *Mosmo^-/-^* embryos. Instead of the predominant TGA phenotype seen in untreated *Mosmo^-/-^* embryos, SMO inhibition shifted the phenotype to DORV, an overlapping conotruncal malformation also associated with defective ventriculoarterial alignment but of reduced severity compared to TGA (**Figures 5A-C** **and Table S4**). Vismodegib treatment in *Mosmo^-/-^* embryos also had a corrective effect on the position of the heart, reducing the incidence of mesocardia, dextrocardia, and right aortic arch (RAA). Notably, these defects are all reflective of improper left-right cardiac morphogenesis (**Figures 5A-B** **and Table S4**). Perhaps the most compelling evidence that vismodegib treatment improved *Mosmo^-/-^* heart development and function came from the analysis of embryo survival. The number of *Mosmo*^-/-^ embryos that survived to e14.5 was 16-fold higher in vismodegib-treated litters compared to vehicle-treated litters (**Figure 5D** **and Table S5**). We speculate that the incomplete rescue of CHDs may be due to the challenges of delivering the correct dose of vismodegib at the correct time to impact the multiple (temporally different) points in development when Hh signaling is required for heart development.

### Neural patterning in *Mosmo^-/-^* embryos is resistant to SMO inhibitors

The loss of MOSMO did not cause defects in all tissues that are known to require SHH for their patterning. Dorsal-ventral patterning of the developing spinal cord is coordinated by a gradient of SHH that is secreted initially from the notochord and later from the floor plate. A large body of literature has shown that neural progenitors adopt different cell fates depending on the magnitude of their exposure to SHH. Increasing the concentration of SHH or the duration of SHH exposure results in an expansion of ventral neural cell fates (Dessaud et al., 2007). Unexpectedly, staining with a panel of cell-type specific markers revealed normal neural tube patterning in *Mosmo*^-/-^ embryos (**Figure S6B**). The integrity of neural patterning was maintained despite the fact that the loss of MOSMO resulted in markedly elevated levels of ciliary SMO along the entire dorsal-ventral axis of the developing spinal cord (**Figure S6C**). In wild-type embryos, SMO accumulates in the cilia of only the ventral-most progenitor cells (floor plate and p3 progenitors) of the developing spinal cord, cells that are exposed to the highest concentrations of SHH (Kong et al., 2015). However, ciliary SMO levels were elevated in all progenitor domains in *Mosmo*^-/-^ embryos, even those distant from the SHH source at the floor plate (**Figures S6C**).

While the patterning of ventral spinal progenitors was indistinguishable in control (*Mosmo^+/+^* and *Mosmo^+/-^*) and *Mosmo*^-/-^ embryos, a dramatic difference was uncovered when these embryos were exposed to vismodegib. Similar to previous experiments, pregnant dams from *Mosmo^+/-^* x *Mosmo^+/-^* crosses were exposed to vismodegib from e7.75-e11.25 and then collected at e11.5 for spinal cord analysis. In control (*Mosmo^+/+^* and *Mosmo^+/-^*) embryos, vismodegib caused the loss of ciliary SMO (**Figure 6A**) and the loss of the ventral-most progenitor domains (OLIG2+ and NKX2.2+ neural progenitors) that require SHH for their specification (**Figure 6B**). These same ventral cell types are also lost in *Smo*^-/-^ and *Shh*^-/-^ embryos, confirming that vismodegib mimics the genetic disruption of Hh signaling (Chiang et al., 1996; Wijgerde et al., 2002). In contrast, *Mosmo*^-/-^ embryos from the same litter (exposed to the same vismodegib regimen), maintained OLIG2+ and NKX2.2+ progenitors (**Figure 6B-D**). Although the OLIG2+ and NKX2.2+ progenitors were present, they were shifted to more ventral positions within the spinal cord in vismodegib-treated *Mosmo^-/-^* embryos, consistent with partial SMO inhibition. Thus, the loss of a single gene, *Mosmo*, can influence the impact of an established teratogen (vismodegib) on neural tube patterning, likely by increasing the abundance of SMO, the protein target of the teratogen.

**Figure 6:**
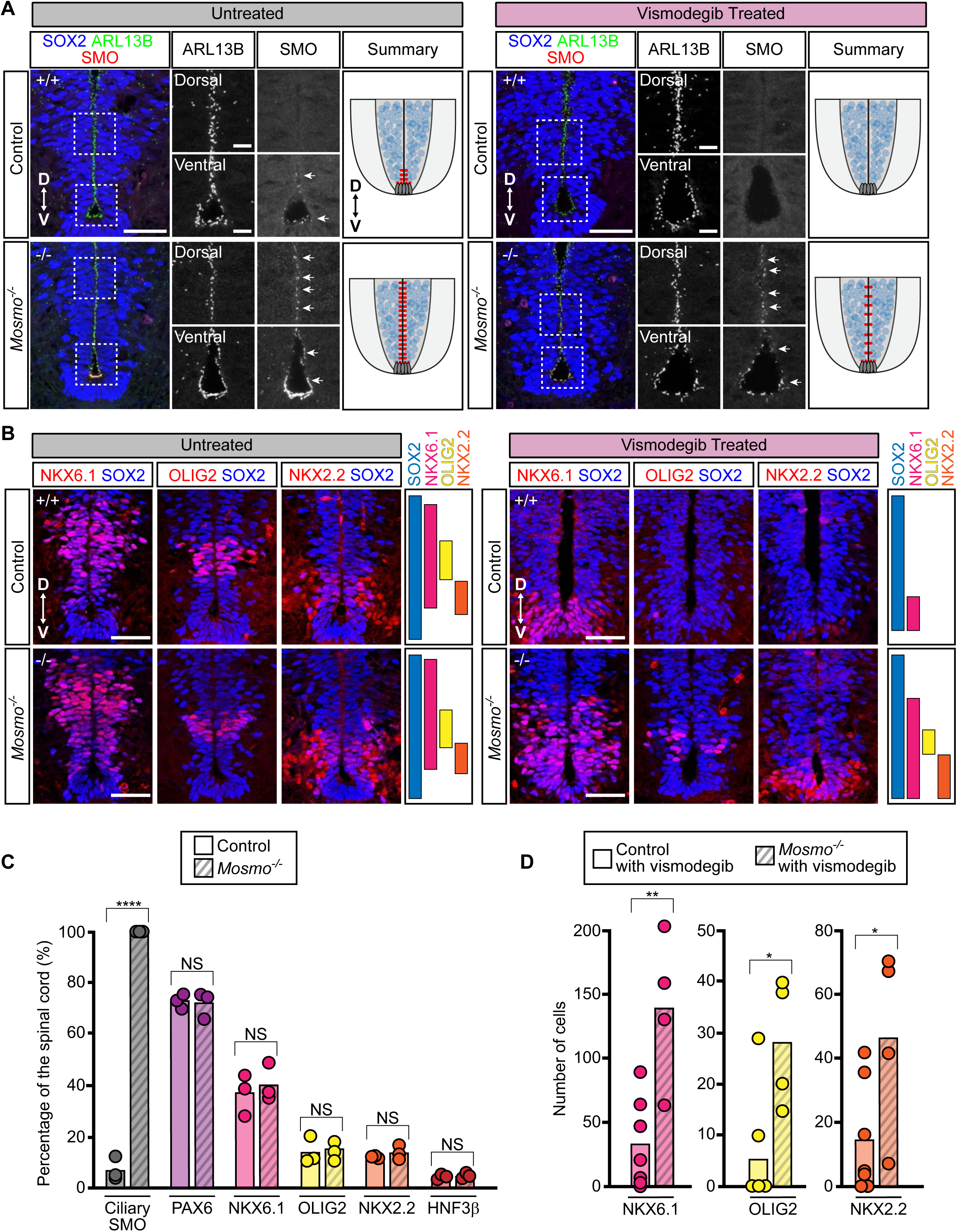
The developing spinal cords of *Mosmo^-/-^* embryos are resistant to SMO inhibition. Neural tube patterning was assessed using confocal fluorescence microscopy to image markers that define neural progenitor populations in sections of the ventral spinal cord from e11.5 control (*Mosmo^+/+^* and *Mosmo^+/-^*) and *Mosmo^-/-^* embryos treated with or without vismodegib. **(A)** Immunofluorescence (IF) was used to evaluate SMO abundance (red) in primary cilia (ARL13B, green) within neural progenitors (SOX2, blue) of untreated e11.5 embryos (left) and vismodegib treated embryos (right). White arrows point to regions where SMO is seen in cilia and red hatch marks in the diagrams denote the regions of the developing neural tube where ciliary SMO is observed. Scale bars, 100 µm in merged panels and 50 µm in zoomed displays. **(B)** IF was used to assess the abundance and distribution of NKX6.1+, OLIG2+, and NKX2.2+ (red) neural progenitor cells (SOX2, blue) of untreated e11.5 embryos (left) and vismodegib treated embryos (right). Images represent three serial sections taken from a single representative embryo. Scale bars are 50 µm. **(C)** Summary of the dorsal-ventral distribution of ciliary SMO, PAX6, NKX6.1, OLIG2, NKX2.2, and HNF3β from untreated e10.5 control (*Mosmo^+/+^* and *Mosmo^+/-^*) and *Mosmo^-/-^* embryos. See **Figure S6B and S6C** for representative images. **(D)** Quantification of NKX6.1+, OLIG2+, and NKX2.2+ spinal neural progenitors from vismodegib treated e11.5 control and *Mosmo^-/-^* embryos. (C-D) Each point represents one embryo. Bars represent the mean and the statistical analysis between the two groups was determined using an unpaired t-test. not-significant (ns) *p*-value > 0.05, \**p*-value ≤ 0.05, \*\**p*-value ≤ 0.01, ****p value ≤ 0.0001.

## DISCUSSION

The MMM complex composed of MOSMO, MEGF8, and MGRN1 anchors a signaling pathway that regulates the sensitivity of target cells to Hh morphogens (**Figure 7**). All three components of the MMM complex were originally identified as attenuators of Hh signaling in our genome-wide CRISPR screens (Pusapati et al., 2018). The phenotypic similarities between *Mosmo^-/-^*, *Megf8^-/-^*, and *Mgrn1^-/-^*;*Rnf157^-/-^* cells and mouse embryos supports the model that the MMM proteins function in the same pathway. We previously found that MGRN1 interacts with MEGF8, forming a membrane-tethered ubiquitination complex that targets SMO for degradation (Kong et al., 2020). Here, we report that MOSMO interacts with MEGF8 to facilitate its accumulation at the cell surface. This mechanism is reminiscent of how tetraspanins like CD81 promote cell-surface expression of the B-cell co-receptor CD19 (Shoham et al., 2003). Since MOSMO was a previously unannotated and unstudied protein, we ablated *Mosmo* in the mouse and discovered that it is essential for embryonic development. In the absence of MOSMO, SMO is enriched in the primary cilia of all tissues, rendering cells hypersensitive to endogenous SHH. Consequently, *Mosmo^-/-^* embryos suffer from severe developmental defects including heterotaxy, skeletal abnormalities, limb patterning defects and congenital heart defects (CHDs).

**Figure 7:**
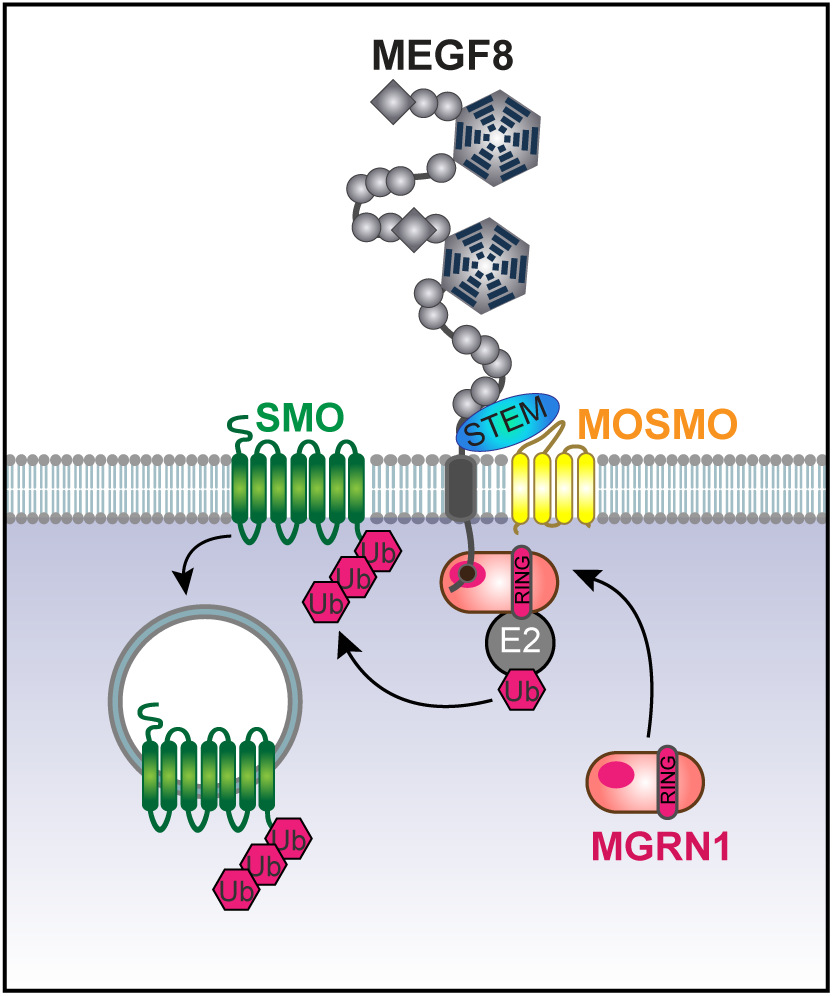
The regulation of cell surface Smoothened (SMO) by the MMM complex. MOSMO binds to the Stem domain of MEGF8. Together, MOSMO, MEGF8, and MGRN1 form a membrane-tethered E3 ligase complex (the MMM complex) that regulates a target cell’s sensitivity to Hh ligands by regulating levels of SMO at the cell surface and primary cilium.

### The MMM complex shapes the Hh signaling gradient

*Mosmo* is widely expressed in the mouse embryo (**Figure S1**) and MOSMO deficiency results in a dramatic increase in ciliary SMO in all tissues we examined (**Figure 2D and 2E**). However, *Mosmo^-/-^* embryos did not show indiscriminate Hh signaling activation in all tissues. The pattern of elevation in Hh signaling activity due to loss of *Mosmo* is very different from the pattern seen in embryos lacking the well-studied Hh negative regulators PTCH1 and SUFU (Cooper et al., 2005; Goodrich et al., 1997; Svärd et al., 2006). While Hh signaling is fully activated in most tissues in *Ptch1^-/-^* and *Sufu^-/-^* embryos, Hh signaling is amplified selectively in SHH-exposed tissues in *Mosmo^-/-^* embryos. We propose that the purpose of the MMM complex is to attenuate the gradient of Hh signaling strength in tissues, rather than to suppress basal signaling activity.

The loss of *Mosmo* clearly has tissue-specific effects. Although Hh signaling is required for the patterning of many tissues, development of the limbs, heart, and skeleton were more severely affected by this elevation in pathway activity than the ventral spinal cord. These differences may reflect whether patterning in a tissue depends on transcriptional de-repression or activation. The patterning of the limb bud is driven by the de-repression of downstream target genes due to a loss of GLI transcriptional repressors (GLIR) (Litingtung et al., 2002; te Welscher et al., 2002). In contrast, the patterning of the ventral spinal cord is primarily driven by the activation of downstream target genes by GLI transcriptional activators (GLIA) (Stamataki et al., 2005). We speculate that loss of the MMM complex potentiates Hh signaling by reducing GLIR levels, rather than by elevating GLIA (Kong et al., 2019; Niewiadomski et al., 2014). In support of this notion, *Mosmo^-/-^* mouse embryos have some of the same phenotypes as *Gli3^-/-^* embryos. GLI3 is proteolytically processed to generate GLI3R, the predominant transcriptional repressor in Hh signaling. As seen in *Mosmo^-/-^* embryos, a loss of GLI3 results in polydactyly (Hui and Joyner, 1993; Johnson, 1967), but no changes in the patterning of the ventral spinal cord (Persson et al., 2002). Overall, our results suggest that limb, heart and skeleton development are particularly susceptible to subtle changes in Hh signaling strength caused by either genetic perturbations or environmental exposures to teratogens like vismodegib. This heightened sensitivity could underlie the sporadic nature of CHDs and contribute to its variable penetrance and expressivity.

### The role of the MMM complex in left-right patterning

Left-right patterning defects, which manifest as heterotaxy with randomization of visceral organ situs, are frequently associated with severe CHDs in humans, suggesting that the signals that specify the left/right body axis also play a role in regulating heart development (Li et al., 2015; Lin et al., 2014). Left-right patterning defects and complex heart malformations are prominent phenotypes common to all the MMM mutant mouse lines (Cota et al., 2006; Zhang et al., 2009). While vismodegib treatment was able to fully rescue the *Mosmo^-/-^* polydactyly phenotype (**Figure 4**), demonstrating that digit duplication is a product of elevated Hh signaling, it failed to fully rescue heterotaxy phenotypes (**Figure 5B**). With regard to the heart, vismodegib partially rescued cardiac *situs* and improved outflow tract malalignment defects, with DORV seen instead of TGA. This finding is intriguing since there is an ongoing debate about whether DORV and TGA are related phenotypes arising from a disturbance in left-right patterning. Our findings suggest that these two phenotypes are indeed developmentally related and may be influenced by changes in Hh signaling strength.

Our failure to rescue heterotaxy phenotypes with vismodegib could be because (1) the MMM complex regulates receptors involved in other signaling pathways or (2) we did not deliver vismodegib during the correct time window in development. Interestingly, a study recently found that the conditional deletion of *Megf8* in all known cardiac cell lineages did not reproduce the heart defects observed in the global *Megf8* knock-out (Wang et al., 2020). These data suggest that Megf8 is required for cardiac development at a time point earlier than cardiac organogenesis and supports the possibility that the heart defects seen in MMM mutant mice are a consequence of disrupted left-right patterning earlier in development. Further studies are needed to identify both the cell types and the critical time periods that are relevant to the function of the MMM complex during development of various tissues.

### Mutations and teratogens converge on Hh signaling to determine the penetrance of birth defects

A central principle of teratology holds that susceptibility to birth defects depends on interactions between the genotype of the embryo and environmental exposures (Finnell, 1999). However, the molecular mechanisms that underlie these gene-teratogen interactions remain largely unknown. Our work on the MMM complex provides one molecular mechanism that explains how mutations and small molecules can interact to influence birth defect outcomes by their combined effects on Hh signaling. Hh ligands function as classical morphogens that direct developmental processes in a manner dependent on the strength of signal in target cells. This simple and elegant mechanism for regulating the patterning and morphogenesis of target tissues, however, also leaves the development of these tissues vulnerable to even small shifts in signaling strength.

Mutations in MMM complex genes cause elevated sensitivity of cells to Hh ligands by increasing the abundance of SMO on the cell surface. Even between embryos in the <u>same litter</u>, *Mosmo* mutations have a profound influence on the teratogenicity of vismodegib, a direct small molecule SMO antagonist. Vismodegib can cause a variety of structural birth defects: neural tube patterning errors, oligodactyly, and cardiac outflow tract abnormalities like PTA (**Figure 4C and 5B**). Remarkably, the developing neural tube and cardiac outflow tracts of *Mosmo^-/-^* embryos are resistant to vismodegib exposure compared to wild-type embryos (**Figure 5 and 6**), likely because the elevated SMO abundance protects these embryos from its teratogenic effects. In the limb and the heart, vismodegib has an even more striking effect on *Mosmo^-/-^* embryos: it rescues structural birth defect phenotypes and improves overall embryo survival, likely by reducing SMO activity to levels that allow normal development. We propose that total SMO activity in target cells, influenced by both SMO protein abundance and exposure to a SMO antagonist, determines birth defect outcomes. Elevated SMO abundance in MMM-mutant embryos can be overcome by reducing SMO activity with vismodegib. Conversely, the reduction in SMO activity caused by vismodegib can be overcome by increasing SMO protein abundance. Thus, gene-environment interactions can arise when genetic factors change the abundance of the protein target of a teratogen.

Prior studies have provided evidence that mutations and environmental exposures can influence development of the face and brain by their combined effects on Hh signaling. Ethanol and the pesticide component piperonyl butoxide (PBO) are exogenous agents that suppress Hh signaling. Early embryonic exposure to high concentrations of ethanol and PBO can cause craniofacial abnormalities and holoprosencephaly (an incomplete division of the forebrain, HPE), phenotypes associated with reduced Hh signaling activity (Ahlgren et al., 2002; Everson et al., 2019; Wang et al., 2012). Embryos exposed to low concentrations of ethanol and PBO develop normally. However, even at low concentrations the teratogenic effects of these agents begin to emerge when they are administered to mice carrying mutations in Hh pathway components (*Shh^+/-^*, *Gli2^+/-^,* or *Cdon^-/-^* embryos) (Everson et al., 2019; Hong and Krauss, 2017; Kietzman et al., 2014).

In summary, the graded, dose-dependent influence of Hh signaling on developmental patterning and morphogenesis explains how gene-teratogen interactions can conspire to modulate the penetrance and expressivity of birth defects by tuning the strength of Hh signaling. A provocative corollary that follows from this idea is that it may be possible to rescue structural birth defect phenotypes by using small molecules (e.g. SMO agonists or antagonists) to re-calibrate Hh signaling strength to the optimal levels required to support normal development. Since these drugs are teratogens themselves, they would need to be delivered during defined time periods in development, at precise doses and to embryos of defined genotypes.

## ACKNOWLEDGEMENTS

We thank Jill Helms, Xue Yuan, and Bo Liu for their assistance with the skeletal stains; Alex Joyner for fruitful discussions, Derek Silvius and Janet Peters for genotyping; and the McLaughlin Research Institute Animal Resource staff for animal care. RR was supported by a grant from the National Institutes of Health (GM118082), GVP by a postdoctoral fellowship from the American Heart Association (14POST20370057), and JHK by a postdoctoral fellowship from the American Heart Association (19POST34380734) and a K99/R00 award from the NIH (GM13251801). CWL, CBY, SH, and GCG were supported by grants from the NIH (HL142788 and HL132024) and the DOD (W81XWH-15-1-0649 and W81XWH-16-1-0613).

## AUTHOR CONTRIBUTIONS

Conceptualization, JHK, GVP, RR, CWL, TMG, JFB; Methodology, JHK, CBY, TMG, JFB; Formal analysis, LA, JHK, CBY, JFB, FB; Investigation, JHK, CBY, GVP, FHE, CBP, SH, BBP, GCG, TMG; Resources, RR, CWL, TMG; Writing--Original Draft, JHK, RR, GVP, JFB, FB; Writing-Revision and Editing, JHK, RR, CWL, TMG, GVP, CBY, CBP; Visualization, JHK, GVP, CBY, JFB; Supervision, RR, CWL, TMG, LA; Project Administration, JHK; Funding Acquisition, RR, CWL, TMG, LA.

## STAR METHODS

### RESOURCE AVAILABILITY

#### Lead Contact

Further information and requests for resources and reagents should be directed to and will be fulfilled by the Lead Contact, Rajat Rohatgi.

#### Materials Availability

All unique/stable reagents generated in this study are available from the Lead Contact with a completed Materials Transfer Agreement.

#### Data and Code Availability

The published article contains all datasets generated and analyzed during this study.

### EXPERIMENTAL MODEL AND SUBJECT DETAILS

#### NIH/3T3 and HEK293T cell culture

Flp-In^TM^-3T3 (referred to as “NIH/3T3” cells throughout the text) and HEK293T cell lines were purchased from Thermo Fisher Scientific and the American Type Culture Collection (ATCC), respectively. Information on the gender of the cell lines is not available. As previously described (Kong et al., 2020), NIH/3T3 and HEK293T cells were cultured in Complete Medium: Dulbecco’s Modified Eagle Medium (DMEM) containing high glucose (Thermo Fisher Scientific, Gibco) supplemented with 10% fetal bovine serum (FBS) (MilliporeSigma), 2 mM L-Glutamine (Gemini Bio-Products), 1 mM sodium pyruvate (Thermo Fisher Scientific, Gibco), 1x MEM non-essential amino acids solution (Thermo Fisher Scientific, Gibco), and penicillin (40 U/ml) and streptomycin (40 µg/ml) (Gemini Bio-Products). The NIH/3T3 and HEK293T cells were rinsed once with sterile PBS and then passaged using 0.05% Trypsin/EDTA (Gemini Bio-Products). Cells were housed at 37 °C in a humidified atmosphere containing 5% CO_2_. Cell lines and derivatives were free of mycoplasma contamination as determined by PCR using the Universal Mycoplasma Detection Kit (ATCC).

#### Generation of primary mouse embryonic fibroblasts

Primary mouse embryonic fibroblasts (pMEFs) were generated using a modified published protocol (Durkin et al., 2013). Briefly, e13.5 embryos were harvested from *Mosmo^+/-^* x *Mosmo^+/-^* crosses and rinsed thoroughly with sterile PBS. Using forceps, the head and internal organs were removed. The embryos were then separated into individual dishes and the tissue was physically minced into small bits in 0.25% Trypsin/EDTA (Thermo Fisher Scientific, Gibco) using a sterile razor blade. Using initially a 5ml serological pipette and later a P1000 pipette tip, the minced tissue was pipetted up and down several times to further break up the tissue, and the dishes were placed in a 37 °C tissue culture incubator for 10-15 minutes. If there were still large tissue pieces present, the minced tissue was pipetted further and the dish was placed in the incubator for an additional 5-10 minutes. The trypsin was then deactivated using Complete Medium (containing 10% FBS). The cells were then centrifuged, resuspended in fresh Complete Medium, and plated. Each clonal cell line represents pMEFs generated from a single embryo. Analysis of *Mosmo^-/-^* pMEFs was always performed with pMEFs prepared from *Mosmo^+/+^* and *Mosmo^+/-^* littermate controls. The gender of the embryos were not determined. Cells were housed at 37 °C in a humidified atmosphere containing 5% CO_2_.

#### Hh signaling assays in NIH/3T3 cells and primary fibroblasts

Hh signaling assays were performed as previously described (Kong et al., 2020). Briefly, NIH/3T3 cells and pMEFs were first grown to confluence in Complete Medium (containing 10% FBS) and then ciliated by changing to Low Serum Medium (Complete Medium containing 0.5% FBS) overnight. NIH/3T3 cells were treated with either no SHH, a low concentration of SHH (1 nM), or a high concentration of SHH (25 nM) prepared in Low Serum Medium. SHH treatment durations varied based on application: 12 hours prior to fixation for NIH/3T3 immunofluorescence assays, 24 hours prior to lysis for NIH/3T3 Western blot assays or NIH/3T3 RNA extraction, and 48 hours prior to pMEF experimentation (immunofluorescence, Western blot, and RNA extraction).

Hh signaling activity was measured using real-time quantitative reverse transcription PCR (qRT-PCR). RNA was extracted from NIH/3T3 cells and pMEFs using TRIzol reagent (Thermo Fisher Scientific, Invitrogen) as previously described (Rio et al., 2010). Equal amounts of RNA were used as a template for cDNA synthesis using the iScript Reverse Transcription Supermix (Bio-Rad Laboratories). qRT-PCR for m*Gli1* and m*Gapdh* was performed on a QuantStudio 5 Real-Time PCR System (Thermo Fisher Scientific) with the following custom designed primers: m*Gli1* (Fwd 5’-CCAAGCCAACTTTATGTCAGGG-3’ and Rev 5’- AGCCCGCTTCTTTGTTAATTTGA-3’) and m*Gapdh* (Fwd 5’-AGTGGCAAAGTGGAGATT-3’ and Rev 5’-GTGGAGTCATACTGGAACA-3’). For all qRT-PCR experiments, *Gli1* transcript levels were calculated relative to *Gapdh* and reported as a fold change across conditions using the comparative C_T_ method (ΔΔC_T_ method).

#### Generation of knockout cell lines

Clonal *Mosmo^-/-^*, *Megf8^-/-^*, *Mgrn1^-/-^* and *Mgrn1^-/-^*;*Rnf157^-/-^* NIH/3T3 lines were previously generated and validated (Kong et al., 2020; Pusapati et al., 2018). Clonal double knockout *Mosmo^-/-^;Megf8^-/-^* NIH/3T3 cell lines were generated using the dual sgRNA strategy to target *Megf8* in *Mosmo^-/-^* NIH/3T3 cells as previously described (Pusapati et al., 2018). Briefly, sgRNAs targeting *Megf8* (5’-TGCCTTCTCTGCCCGAATTG-3’ and 5’- ATAACTTCTCCACGAACACC-3’) were cloned into pSpCas9(BB)-2A-GFP (Addgene) (Ran et al., 2013) and pSpCas9(BB)-2A-mCherry and transfected into NIH/3T3 cells using X-tremeGENE 9 DNA transfection reagent (Roche Molecular Systems). Five days post transfection, GFP and mCherry double positive single cells were sorted into a 96-well plate using a FACSAria II at the Stanford Shared FACS Facility. To detect the GFP, a 488 nm (blue) laser was used with a 530/30 filter and B530 detector. To detect the mCherry, a 561 nm (yellow) laser was used with a 616/23 filter and G616 detector. Clonal lines were screened by PCR (Forward Primer: 5’-CCTCATGCTTGTCCCTTGTT-3’ and Reverse Primer: 5’-GGAGTGTGGGCAAGAAGAAG-3’) to detect excision of the genomic DNA (196 bp) between the two sgRNA cut sites. Knockout of MEGF8 was further confirmed by immunoblotting (**Figure S4D**).

#### Generation of stable cell lines expressing transgenes

*Mosmo^-/-^* NIH/3T3 cells with stable addback of tagged MOSMO (featured in **Figures S4A and S4B**) were generated using the lentiviral expression system as previously described (Kong et al., 2020). Briefly, to generate lentivirus, four million HEK293T cells were seeded onto a 10 cm plate and 24 hours later transfected with 1 µg pMD2.G (Addgene), 5 µg psPAX2 (Addgene), and 6 µg of the *Mosmo-1D4* pLenti CMV Puro DEST construct using 36 µl of 1mg/ml polyethylenimine (PEI) (Polysciences). Approximately 48 hours post transfection, the lentivirus was harvested and filtered through a 0.45 µm filter. 2 ml of the filtered lentivirus solution was mixed with 2 ml of Complete Medium containing 16 µg/mL polybrene (MilliporeSigma). The diluted virus was then added to NIH/3T3 cells seeded on 6-well plates. Approximately 48 hours post infection, cells were split and selected with puromycin (2 µg/ml) for 5-7 days or until all the cells on the control plate were dead.

#### Established Mouse Lines

All mouse studies were conducted using animal study protocols approved by the Institutional Animal Care and Use Committee (IACUC) of Stanford University, the University of Pittsburgh, and the McLaughlin Research Institute for Biomedical Sciences. *Gli1^tm2Alj^* mice (referred to in the paper as *Gli1^lacz^*) (MGI:2449767) and *Megf8^C193R/C193R^* mice (referred to in the paper as *Megf8^m/m^*) (MGI:3722325) have been described previously (Bai et al., 2002; Zhang et al., 2009). *Gli1^lacz^* mice were genotyped using the following primers: Fwd (common) 5’-GGGATCTGTGCCTGAAAC TG-3’, Rev (wild-type) 5’-AGGTGAGACGACTGCCAAGT-3’, and Rev (mutant) 5’-TCTGCCAGTTTGAGGGGACGAC-3’.

#### Generation and genotyping of *Mosmo^-/-^* mutant mice

*Mosmo^+/-^* mice were generated by the Stanford Transgenic Knockout and Tumor Model Center using CRISPR/Cas9 mediated genome editing. In brief, four gRNAs were designed to delete exon 1 of *Mosmo*, to create a downstream reading frame shift in exon 2 and exon 3, and ultimately remove its function (**Figure S2A**). The four guide RNAs (5’-CCGGCGCGCGGTTTCGCTTC-3’, 5’-CGCGGTTTCGCTTCCGGGTG-3’, 5’-CCCCGGGTCGGCGATCCCGA-3’, and 5’-CCCTCGGGATCGCCGACCCG-3’) and CAS9 protein were obtained from Integrated DNA Technologies. A ribonucleoprotein (RNP) injection mix was prepared, consisting of gRNAs (15 ng/ul) and CAS9 protein (30 ng/ul), and introduced into C57BL/6 mouse zygotes via pronuclear microinjection. DNA from the 30 pups born was amplified using primers flanking *Mosmo* exon 1 and sequenced. One male founder that had a 386 bp deletion, including all of exon 1, was backcrossed with C57BL/6 females for two generations, then heterozygotes were intercrossed. *Mosmo* knockout mice were genotyped using two sets of primers to detect the wild-type or KO allele: Fwd (wild-type set 1) 5’-GATAAACTGACCATCATCTCAGGATG-3’, Rev (wild-type set 1) 5’-ACTTCAAAGGGGAAAGGGGGAG-3’, Fwd (wild-type set 2) 5’-GGGCGATGGATAAACTGACC-3’, Rev (wild-type set 2) 5’-CGCCTCTTTCTTGAGGACAC-3’, Fwd (mutant set 1) 5’-CCAGTTCCTTCCCATTGCATCT-3’, Rev (mutant set 1) 5’-GCAGTTCAAATACAAGACCGTTCC-3’, Fwd (mutant set 2) 5’-CCGAGAGCTGGGATTCGTAG-3’, and Rev (mutant set 2) 5’-CCACAGACACTTCAAAGGGGA-3’ (**Figure S2B**).

#### scRNA-seq analysis

We utilized single-cell transcriptome data from the *MouseGastrulationData* R package (Griffiths J, Lun A, 2020), for mouse gastrulation at e7.5 and e8.5. We used Seurat (version 3.2.3) to analyze the scRNA-seq data (Pijuan-Sala et al., 2019). Our workflow for processing the scRNA-seq data involves data pre-processing, centered log ratio transformation across features, scaling with a linear model, dimensionality reduction, and visualization. The annotation of cell types was based on metadata labels included in the data. Cells were plotted based on their euclidean coordinates after a UMAP dimensionality reduction, and *Mosmo* expression is given in normalized read counts.

#### Molecular modeling

The 184 amino acid Stem domain from human MEGF8 comprises amino acids 2463-2647, and is tightly sandwiched between an EGFL module and the hydrophobic transmembrane helix. The Hhpred program in the MPI bioinformatics toolkit (Gabler et al., 2020) was used to define the boundaries and β-sheet nature of this elusive domain by sequence and structural profile matching to MEGF8 orthologs, and Attractin (ATRN) and Attractin-like 1 (ATRNL1) paralogs. The structure of the isolated MEGF8 Stem domain (where a 32 residue, disordered Pro and Gly rich insert, amino acids 2530-2562, relative to a compact ATRN loop, was replaced by a Gly) was predicted and modeled by trRosetta based on its de novo folding algorithm in a template-free fashion, guided by deep learning-derived restraints of residue distances and orientations (Yang et al., 2020). The confidence of the predicted model is high with an estimated TM-score of 0.567. Similar folds present in other PDB structures were revealed by PDBeFOLD searches (Krissinel and Henrick, 2004) with the trRosetta-derived models of MEGF8 and related ATRN and ATRNL1 Stem domains. Residue conservation profiles were mapped to the Stem domain structure with the ConSurf program (Ashkenazy et al., 2016).

### METHOD DETAILS

#### Constructs

*MEGF8*-1D4, *MEGF8*ΔN-1D4, *MEGF8*ΔC-1D4, *Mgrn1*-3xFLAG, *Mgrn1^Mut1^*-3xFLAG, *Smo*-EGFP, and *Mosmo*-1D4 were previously described (Kong et al., 2020; Pusapati et al., 2018). *Mosmo*-3xHA was synthesized as a gBlock (Integrated DNA Technologies) and used as a template for the PCR amplification step. To generate MEGF8ΔN^Stem^, *MEGF8* (NM_001410.3) nucleotide sequence coding for amino acids 2313-2778 was PCR amplified using *MEGF8* full length as a template. All constructs were cloned initially into pENTR2B plasmid (Thermo Fisher Scientific, Invitrogen) and then transferred into pEF5/FRT/V5-DEST (Thermo Fisher Scientific, Invitrogen) or pLenti CMV PURO DEST (Campeau et al., 2009) using Gateway recombination methods (Thermo Fisher Scientific, Invitrogen).

#### Reagents and antibodies

Recombinant SHH was expressed in bacteria and purified in the lab as previously described (Bishop et al., 2009). Briefly, His-tagged SHH-N (C24II followed by human SHH amino acids 25-193) was expressed in *Escherichia coli* (BL21 strain; Rosetta2 (DE3)pLysS). Cells were lysed in 10 mM Phosphate Buffer pH 7.5, 500 mM NaCl, 1 mM 2-mercaptoethanol, 1 mM PMSF, and 1x protease inhibitor cocktail, followed by centrifugation at 20,000x*g* for 30 minutes at 4°C. Clarified samples were incubated with Ni-NTA resin (Qiagen) for 1 h at 4°C. The resin was washed with 20 column volumes of wash buffer A (lysis buffer without protease inhibitors), followed by wash buffer B (wash buffer A + 10 mM Imidazole), and bound proteins were eluted with elution buffer (wash buffer A + 250 mM Imidazole). Peak fractions were pooled, concentrated using a 5 kDa cut-off VIVASPIN 15R (Life Technologies), and loaded onto a Superdex 75 gel filtration column (Amersham Biosciences) equilibrated with column buffer (10 mM HEPES pH 7.5, 150 mM NaCl, and 1 mM DTT). The recombinant protein was >98% pure, as assessed from coomassie staining and stored at -80 °C. The selection antibiotic puromycin was purchased from MilliporeSigma. The transfection reagent XtremeGENE 9 was purchased from Roche Molecular Systems and polybrene from MilliporeSigma. Bafilomycin A1 was purchased from Cayman Chemical. Vismodegib and Bortezomib were purchased from LC labs. The following primary antibodies were purchased from the following vendors: mouse anti-1D4 (The University of British Columbia, 1:5000); rat anti-E-cadherin (clone ECCD-2, Zymed, 1:1000); mouse anti-FLAG (clone M2, MilliporeSigma, 1:2000); goat anti-GFP (Rockland Immunochemicals, 1:1000); rabbit anti-GFP (Novus Biologicals, 1:5000); mouse anti-GLI1 (clone L42B10, Cell Signaling, 1:1000); mouse anti-HA (clone 2-2.2.14, Thermo Fisher Scientific, 1:2000); rabbit anti-p38 (Abcam, 1:2000); and rabbit anti-RNF156 (anti-MGRN1, Proteintech, 1:500); mouse anti-ɑ-Tubulin (Clone DM1A, MilliporeSigma, 1:10000); mouse anti-acetylated-Tubulin (MilliporeSigma, 1:10000). The following primary antibodies were generated in the lab or received as a gift: Guinea pig anti-ARL13B (1:1000) (Dorn et al., 2012); rabbit anti-SMO (designed against an intracellular epitope, 1:2000) (Rohatgi et al., 2007); and rabbit anti-MEGF8 (1:2000) (Kong et al., 2020). Hoechst 33342 and secondary antibodies conjugated to horseradish peroxidase (HRP) or Alexa Fluor dyes were obtained from Jackson ImmunoResearch Laboratories and Thermo Fisher Scientific.

#### Immunoprecipitation and Western Blotting

Whole cell extracts from HEK293T and NIH/3T3 cells were prepared in Immunoprecipitation (IP) Lysis Buffer: 50 mM Tris pH 8.0, 150 mM NaCl, 1% NP-40, 0.25% sodium deoxycholate, 1 mM DTT, and 1x SIGMAFAST protease inhibitor cocktail (MilliporeSigma). Cells were lysed for 1 hour on a shaker at 4 °C, supernatants were clarified by centrifugation (20,000 xg for 30 minutes at 4 °C), and 1D4 tagged MOSMO or MEGF8 was captured by a 1D4 antibody (The University of British Columbia) covalently conjugated to Protein A Dynabeads (Thermo Fisher Scientific, Invitrogen). Immunoprecipitates were washed once with IP Wash Buffer A (50 mM Tris at pH 8.0, 150 mM NaCl, 1% NP-40, 0.25% sodium deoxycholate, and 1 mM DTT), once with IP Wash Buffer B (50 mM Tris at pH 8.0, 500 mM NaCl, 0.1% NP-40, 0.25% sodium deoxycholate, and 1 mM DTT), and finally with IP Wash Buffer C (50 mM Tris at pH 8.0, 0.1% NP-40, 0.25% sodium deoxycholate, and 1 mM DTT). Proteins were eluted by resuspending samples in 2x NuPAGE LDS sample buffer (Thermo Fisher Scientific, Invitrogen) supplemented with 100 mM DTT, incubated at 37 °C for 1 hour, and subjected to SDS-PAGE (**Figures 3E and 3G**).

Whole cell extracts were prepared in RIPA lysis buffer: 50 mM Tris at pH 8.0, 150 mM NaCl, 2% NP-40, 0.25% sodium deoxycholate, 0.1% SDS, 0.5 mM TCEP, 10% glycerol, 1x SIGMAFAST protease inhibitor cocktail (MilliporeSigma), and 1x PhosSTOP phosphatase inhibitor cocktail (Roche). The resolved proteins were transferred onto a nitrocellulose membrane (Bio-Rad Laboratories) using a wet electroblotting system (Bio-Rad Laboratories) followed by immunoblotting.

For the preparation of whole embryo extracts, e12.5 embryos were collected and rinsed thoroughly in chilled PBS. Each embryo was then individually submerged in liquid nitrogen and pulverized using a mortar and pestle. The crushed tissue was then lysed in modified RIPA lysis buffer: 50 mM Tris at pH 8.0, 150 mM NaCl, 1% NP-40, 0.25% sodium deoxycholate, 0.1% SDS, 5 mM EDTA, 1 mM sodium fluoride, 1mM sodium orthovanadate, and 1x SIGMAFAST protease inhibitor cocktail (MilliporeSigma). The resolved proteins were then transferred onto a nitrocellulose membrane (Bio-Rad Laboratories) using a wet electroblotting system (Bio-Rad Laboratories) and imunoblotted.

#### Cell surface biotinylation assay

Cell surface levels of MEGF8 (**Figure 3B**) were determined by a biotinylation assay as described previously (Kong et al., 2020). Briefly, wild-type, *Mosmo^-/-^*, and *Megf8^-/-^* NIH/3T3 cells were plated on 10 cm plates in Complete Medium. Once the cells were confluent they were switched to Low Serum Medium for 24 h. On biotinylation day, the cells were removed from the 37 °C incubator and placed on an ice-chilled metal rack in a 4 °C cold room. The medium was removed and cells were quickly washed 3 times with ice-cold DPBS+ buffer (Dulbecco’s PBS supplemented with 0.9 mM CaCl_2_, 0.49 mM MgCl_2_.6H_2_O, 5.6 mM dextrose, and 0.3 mM sodium pyruvate). Biotinylation of cell surface proteins using a non-cell permeable and thiol-cleavable probe was initiated by incubating cells with 0.4 mM EZ-Link Sulfo-NHS-SS-Biotin (Thermo Fisher Scientific) in DPBS+ buffer for 30 min. Unreacted Sulfo-NHS-SS-Biotin was quenched with 50 mM Tris (pH 7.4) for 10 min. Cells were then washed 3 times with a 1x Tris-buffered saline (25 mM Tris at pH 7.4, 137 mM NaCl, and 2.7 mM KCl) and whole cell extracts were prepared in Biotinylation Lysis Buffer A (50 mM Tris at pH 8.0, 150 mM NaCl, 2% NP-40, 0.25% sodium deoxycholate, 1x SIGMAFAST protease inhibitor cocktail (MilliporeSigma), and 1x PhosSTOP phosphatase inhibitor cocktail (Roche). Biotinylated proteins from clarified supernatants were captured on a streptavidin agarose resin (TriLink Biotechnologies), washed once with Biotinylation Lysis Buffer A, once with Biotinylation Wash Buffer A (Biotinylation Lysis Buffer A + 0.5% SDS), once with Biotinylation Wash Buffer B (Biotinylation Wash Buffer A + 150 mM NaCl), and finally once again with Biotinylation Wash Buffer A. Biotinylated proteins captured on streptavidin agarose resin were eluted in 1x NuPAGE-LDS sample buffer (Thermo Fisher Scientific, Invitrogen) containing 100 mM DTT at 37 °C for 1 hour and assayed by immunoblotting for MEGF8.

#### Ubiquitination assay

Ubiquitination assays were performed as previously reported (Kong et al., 2020). Briefly, 8 million HEK293T cells were plated onto a 15 cm plate. 24 hours after plating, the cells were transfected using PEI. 6 ug of each construct was transfected into the HEK293T cells at a DNA:PEI ratio of 1:3. An empty plasmid construct was used as filler DNA to ensure that each plate was transfected with the same amount of DNA. To enrich for ubiquitinated proteins, 36 hours post transfection, cells were pre-treated with 10 µM Bortezomib (a proteasome inhibitor) and 100 nM Bafilomycin A1 (a lysosome inhibitor) for 4 hours. Cells were washed twice with chilled PBS and lysed in Ubiquitination Lysis Buffer A comprised of: 50 mM Tris at pH 8.0, 150 mM NaCl, 2% NP-40, 0.25% sodium deoxycholate, 0.1% SDS, 6M urea, 1 mM DTT, 10 µM Bortezomib, 100 nM Bafilomycin A1, 20 mM N-Ethylmaleimide (NEM, MilliporeSigma), and 1x SIGMAFAST protease inhibitor cocktail (MilliporeSigma). Clarified supernatants were diluted ten-fold with Ubiquitination Lysis Buffer B (Ubiquitination Lysis Buffer A prepared without urea) to adjust the urea concentration to 600 mM. For these assays, we assessed ubiquitination on GFP tagged SMO. Ubiquitinated GFP tagged SMO (**Figures 3C and 3D**) was captured using a GFP binding protein (GBP) covalently conjugated to carboxylic acid decorated Dynabeads (Dynabeads M-270 carboxylic acid, Thermo Fisher Scientific). Immunoprecipitates were washed once with Ubiquitination Wash Buffer A (Ubiquitination Lysis Buffer B + 0.5% SDS), once with Ubiquitination Wash Buffer B (Ubiquitination Wash Buffer A + 1 M NaCl), and finally once again with Ubiquitination Wash Buffer A. Proteins bound to dynabeads were eluted in 2x NuPAGE-LDS sample buffer (Thermo Fisher Scientific, Invitrogen) containing 30 mM DTT at 37 °C for 1 hour and assayed by immunoblotting.

#### Immunofluorescence staining of cells and tissue and image quantifications

Mouse tissue was prepared for immunofluorescence imaging as previously described (Kong et al., 2020). Briefly, mouse embryos of various ages were harvested and fixed in 4% (w/v) paraformaldehyde (PFA) in PBS at 4 °C on a nutator. Fixation time varied depending on the age of the embryo (30 minutes for e8.5-9.5, 1 hour for e10.5-11.5, and 2 hours for e12.5). The embryos were then rinsed thoroughly in chilled PBS. To cryopreserve the tissue, the embryos were transferred to 30% sucrose in 0.1M PB (pH 7.2) and allowed to equilibrate overnight. The embryos were then carefully dissected, then the desired tissues were mounted and frozen into Tissue-Plus OCT (optimal cutting temperature) compound (Thermo Fisher Scientific) and 12-14 µm sections were collected on a Leica CM1800 cryostat. Prior to staining, the tissues were blocked for 1 hour at room temperature in immunofluorescence (IF) Blocking Buffer: 1% normal donkey serum (NDS) and 0.1% Triton-X diluted in PBS. In a humidified chamber, the sections were then incubated with primary antibodies overnight prepared in IF Blocking Buffer at 4 °C, rinsed 3 times in PBST (PBS + 0.1% Triton-X), incubated with secondary antibodies and Hoescht prepared in IF Blocking Buffer for 1 hour at room temperature, rinsed 3 times in PBST, and then mounted in ProLong^TM^ Gold antifade mountant (Thermo Fisher Scientific, Invitrogen).

NIH/3T3 cells and pMEFs were fixed in chilled 4% (w/v) PFA in PBS for 10 minutes and then rinsed thoroughly with chilled PBS. Cells were incubated in IF Blocking Buffer for 30 minutes, primary antibodies for 1 hour, and secondary antibodies for 30 minutes. Coverslips were mounted in ProLong^TM^ Gold antifade mountant (Thermo Fisher Scientific, Invitrogen).

Fluorescent images were acquired on an inverted Leica SP8 confocal microscope equipped with a 63X oil immersion objective (NA 1.4). Z-stacks (∼4 µm sections) were acquired with identical acquisition settings (laser power, gain, offset, frame and image format) within a given experiment. An 4-8X optical zoom was used for imaging cilia to depict representative images. For the quantification of SMO at cilia, images were opened in Fiji (Schindelin et al., 2012) with projections of the maximum fluorescent intensities of z-stacks. Ciliary masks were constructed based on ARL13B images and then applied to corresponding SMO images to measure the fluorescence intensity of SMO at cilia.

#### Vismodegib dosing via oral gavage

Vismodegib treatment was performed as described previously (Heyne et al., 2015). Briefly, *Mosmo^+/-^* x *Mosmo^+/-^* and wild-type x *Gli1^lacz/+^* mouse crosses were set up and monitored daily. Time e0 was defined as midnight prior to visualization of the copulation plug. Female mice were weighed at ∼e0.25 (the morning the plug was visualized) and ∼e7.25. Only mice that gained 1.75 grams over 7 days were deemed “likely pregnant” and treated with either vehicle or vismodegib. For vismodegib treatment, a 3 mg/ml vismodegib solution was prepared in 0.5% methyl cellulose (MilliporeSigma) with 0.2% Tween. Vismodegib (40 mg/kg) was administered via oral gavage every 12 hours (∼7am and 7pm) for a total of three days (e8.25-e10.75) or four days (e7.25-e10.75) (**Figures 4C, 5A-D**, and **Table S4**). Embryos were harvested at e14.5, fixed in 4% (w/v) PFA in PBS for at least 24 hours, and then analyzed.

#### Mouse embryo phenotyping analysis

Mouse embryo phenotyping was performed as described previous (Kong et al., 2020). Briefly, mouse embryos (e14.5) were fixed in 4% (w/v) PFA in PBS for at least 24 hours. Necropsy was performed to determine visceral organ situs (i.e. lung and liver lobation, heart directionality, and positioning of the stomach, spleen, and pancreas) (**Figure 3A, Tables S2 and S4**). For analysis by episcopic confocal microscopy (ECM), the samples were embedded in paraffin and processed as previously described (Liu et al., 2013). Briefly, the tissue block was sectioned using a Leica sledge microtome and serial images of the block face were captured with a Leica confocal microscope. The serial two-dimensional (2D) image stacks generated were then three-dimensionally (3D) reconstructed using Osirix software (Rosset et al., 2004) and digitally resliced in different orientations to aid in the analysis of intracardiac anatomy and the diagnosis of congenital heart defects (Liu et al., 2013) (**Figures 1F and 5A**).

#### Whole-mount skeletal staining

Whole-mount skeletal stains were prepared using a modified published protocol (McLeod, 1980; Rigueur and Lyons, 2014). Briefly, e15.5-e16.5 mouse embryos were harvested and the skin and internal organs were removed to facilitate tissue permeabilization. The embryos were then fixed first in 95% ethanol (EtOH) overnight at room temperature followed by 100% acetone overnight at room temperature. To stain the cartilage, the embryos were incubated overnight at room temperature in 0.03% (w/v) Alcian Blue 8GX (Millipore Sigma) prepared in a solution of 80% EtOH and 20% glacial acetic acid. After visually confirming that the embryos were completely blue, the embryos were destained in a series of EtOH washes (2-3 hours in 100% EtOH, 75% EtOH, 50% EtOH, and 25% EtOH). To stain the bone, the embryos were then incubated overnight at 4 °C in 0.005% (w/v) Alizarin Red S (Millipore Sigma) prepared in 1% potassium hydroxide (KOH). The tissue was then cleared in 0.3% KOH for 1 to 3 days (changing the solution every day). Once the embryos cleared to the desired amount, the 0.3% KOH was replaced with glycerol. The embryos were transitioned through a series of glycerol solutions (20% for 1 day, 50% for 1 day, and then 80% for 1 day). The skeletons were then kept in 80% glycerol for prolonged storage.

#### Whole-mount lung staining and branching analysis

Whole-mount lungs were prepared as previously described (Metzger et al., 2008). Briefly, e12.5 mouse embryos were harvested and the lungs were carefully excised. The lungs were fixed in 4% (w/v) PFA in PBS for 1 hour and then rinsed thoroughly in PBS at room temperature. The lungs were dehydrated in a series of methanol (MeOH) washes: once in 25% MeOH/PBS (v/v), once in 50% MeOH/PBS, once in 75% MeOH/PBS, and twice in 100% MeOH. The dehydrated lungs were then bleached for 15 minutes in 5% H_2_O_2_/MeOH at room temperature and then rehydrated in a series of PBT washes (PBS with 0.1% Tween-20): once in 75% MeOH/PBS, once in 50% MeOH/PBT, once in 25% MeOH/PBT, and thrice in 100% PBT). The lungs were blocked for 1 hour at room temperature in Whole-mount (WM) Blocking Buffer: 5% donkey serum and 0.5% Triton X-100 diluted in PBS. For primary antibody labeling, the lungs were incubated overnight at 4 °C in rat anti-E-cadherin antibody (clone ECCD-2, Zymed) diluted 1:1000 in WM Blocking Buffer. The following day, the lungs were thoroughly rinsed in PBT (8 x 30 minutes). For secondary antibody labeling, the lungs were incubated overnight at 4 °C in biotin-conjugated donkey anti-rat IgG (Jackson ImmunoResearch Laboratories) diluted 1:250 in WM Blocking Buffer. The lungs were then thoroughly rinsed in PBT (8 x 30 minutes), the biotin was visualized using the VECTASTAIN Elite ABC Kit (Vector, PK-16100), and the signal was amplified using a Tyramide Signal Amplification System (Cy3, Perkin Elmer). Stained lungs were mounted in Vectashield with DAPI (Vector) and imaged on a Thunder Imager Model Organism (Leica) (**Figure 1E** **and S3A**).

#### Whole-mount β-galactosidase staining of mouse embryos

Mouse embryos were processed for β-galactosidase staining using a modified published protocol (Nagy et al., 2007). Briefly, embryos were harvested from *Mosmo^+/-^* x *Mosmo^+/-^*;*Gli1^lacZ/+^* mouse crosses and fixed at 4 °C in 4% (w/v) PFA in PBS for varying durations of time depending on their age (e8.5 and e9.5 for 10 minutes, e10.5 for 12 minutes, e11.5 for 15 minutes, and e12.5 for 20 minutes). The embryos were then rinsed thoroughly in PBS and permeabilized for either 2 hours (≤e9.5) or overnight (≥10.5) at 4 °C in Permeabilization Solution: 0.02% sodium deoxycholate and 0.01% NP-40 diluted in PBS. Following permeabilization, the embryos were placed in Staining Solution: 2mM MgCl_2_, 0.02% NP-40, 5 mM potassium ferricyanide, 5 mM potassium ferrocyanide, 0.01% sodium deoxycholate diluted in 0.1M phosphate buffer (pH 7.2). The embryos were stained for 2 hours at 37°C. To remove residual yellow color from the Staining Solution, the embryos were rinsed in Permeabilization Solution (2 x 15 minutes). The embryos were fixed overnight in 4% (w/v) PFA in PBS at 4 °C, rinsed in PBS, and then imaged.

#### In situ hybridization of whole-mount and sectioned tissue

As previously described (Pusapati et al., 2018), to generate a *Mosmo* in situ probe, *Mosmo* specific primers were designed using the program Primer3: forward 5’-acacgtgtgtgctgaaaagc-3’ and reverse 5’-gag<u>attaaccctcactaaaggga</u>tgagcaggtaacccatctcc-3.’ The underlined sequence marks the T3 polymerase binding site that was incorporated into the reverse primer. The *Mosmo* probe was generated using a Digoxigenin (DIG) RNA Labeling Kit (Roche). Briefly, the probe was generated from the in vitro transcription of PCR products amplified from mouse neural progenitor cell cDNA. After overnight hybridization at 65 °C, the signal was visualized using Anti-DIG-alkaline phosphatase (AP) Fab fragments (Roche) and NBT/BCIP (Roche).

### QUANTIFICATION AND STATISTICAL ANALYSIS

Most data was analyzed using GraphPad Prism 9. Violin plots (**Figure 2D** **and S4A**) were created using the “Violin plot (truncated)” appearance function. In Prism 9, the frequency distribution curves of the violin plots are calculated using kernal density estimation. By using the “truncated” violin plot function, the frequency distributions shown are confined within the minimum to maximum values of the data set. On each violin plot, the median (central bold line) and quartiles (adjacent thin lines, representing the first and third quartiles) are labeled. In Prism 9, the statistical significance between two groups was determined using either Mann-Whitney (**Figure 2D**) or an unpaired t-test (**Figure 6C and 6D**) and the significance between three or more groups was determined using the Kruskal-Wallis test (**Figure 4C** **and S4A**). For each of these figures, *p*-values were calculated using Prism 9 and reported in the figure legend using the following key: not-significant (ns) *p*-value > 0.05, \**p*-value ≤ 0.05, \*\**p*-value ≤ 0.01, \*\*\**p*-value ≤ 0.001, and \*\*\*\**p*-value ≤ 0.0001 Additional figure details regarding the *n*-value and statistical test applied were reported in the individual figure legends.

Disruptions in offspring viability due to genotype (**Figure 1A** **and Table S1**) or treatment (**Figure 5D** **and Table S5**) were determined using the chi-square test. Briefly, *Mosmo^+/-^* x *Mosmo^+/-^* crosses were set up, live embryos were collected, and deviation from the expected Mendelian ratio of 1:2:1 was calculated. Not-significant (ns) > 0.05, \*\**p*-value ≤ 0.01, and ***p value ≤ 0.001.

All cell biological and biochemical experiments were performed two to three independent times, with similar results. To validate *Mosmo^-/-^* primary mouse embryonic fibroblasts (pMEFs), 3 independent cell lines were generated (each from a single embryo) and compared against control (*Mosmo^+/+^* and *Mosmo^+/-^*) pMEFs generated from embryos within the same litter (**Figure 2A**). Similarly, 2 whole embryo lysate samples were prepared (each from a single *Mosmo^-/-^* embryo) and compared against control (*Mosmo^+/+^* and *Mosmo^+/-^*) lysates prepared from embryos within the same litter (**Figure 2C**).

---

**Figure S1:**
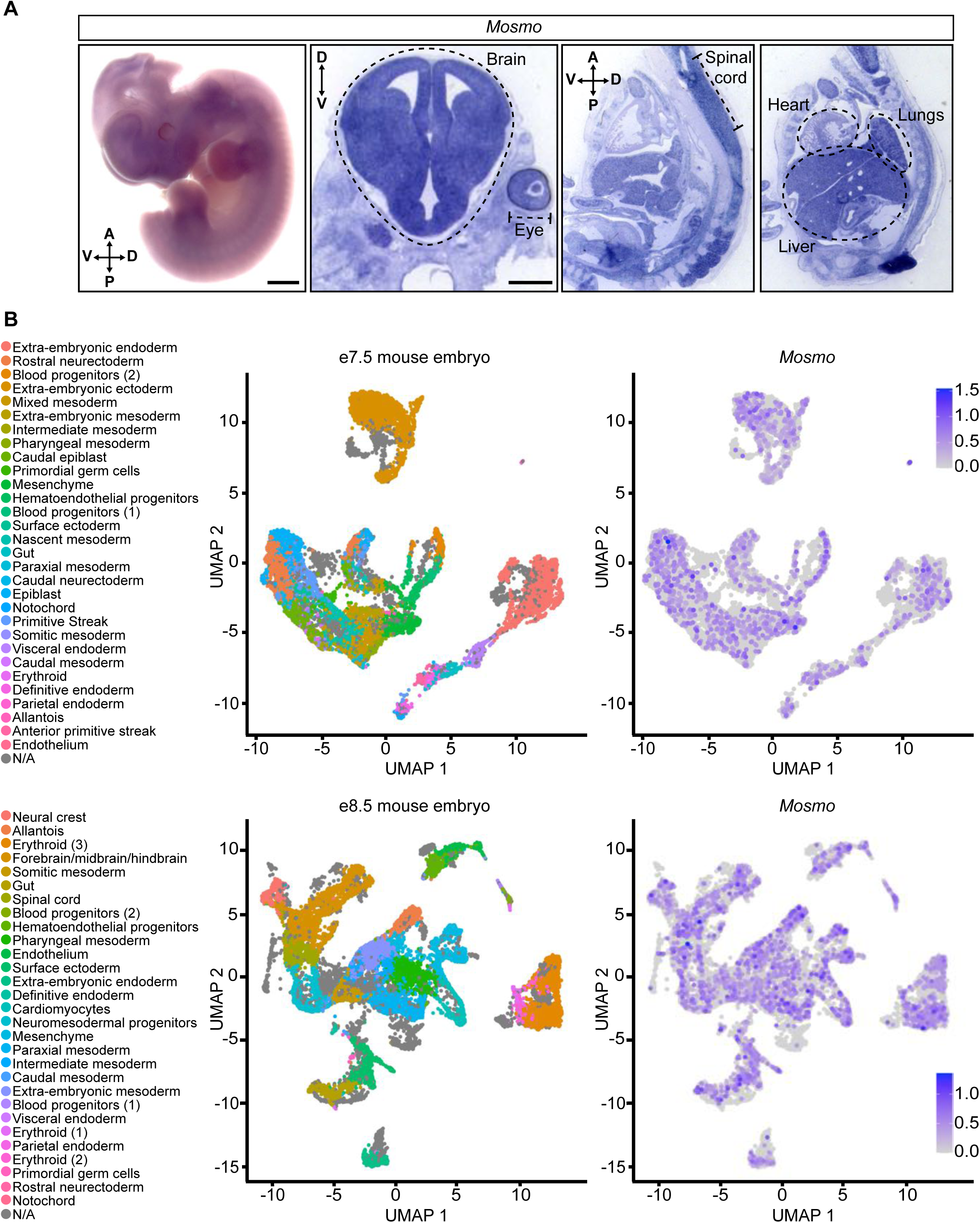
*Mosmo* is widely expressed in the developing embryo. **(A)** Whole-mount *in situ* hybridization was used to assess *Mosmo* expression in whole-mount mouse embryo preparations (e11.5, leftmost panel) or in coronal brain and sagittal whole body sections (e13.5, three right panels). Scale bars are 1 mm. **(B)** UMAP analysis of previously published single-cell RNAseq data (Pijuan-Sala et al., 2019) showing *Mosmo* expression in an abundance of single cells from e7.5 (top) and e8.5 (bottom) embryos.

**Figure S2:**
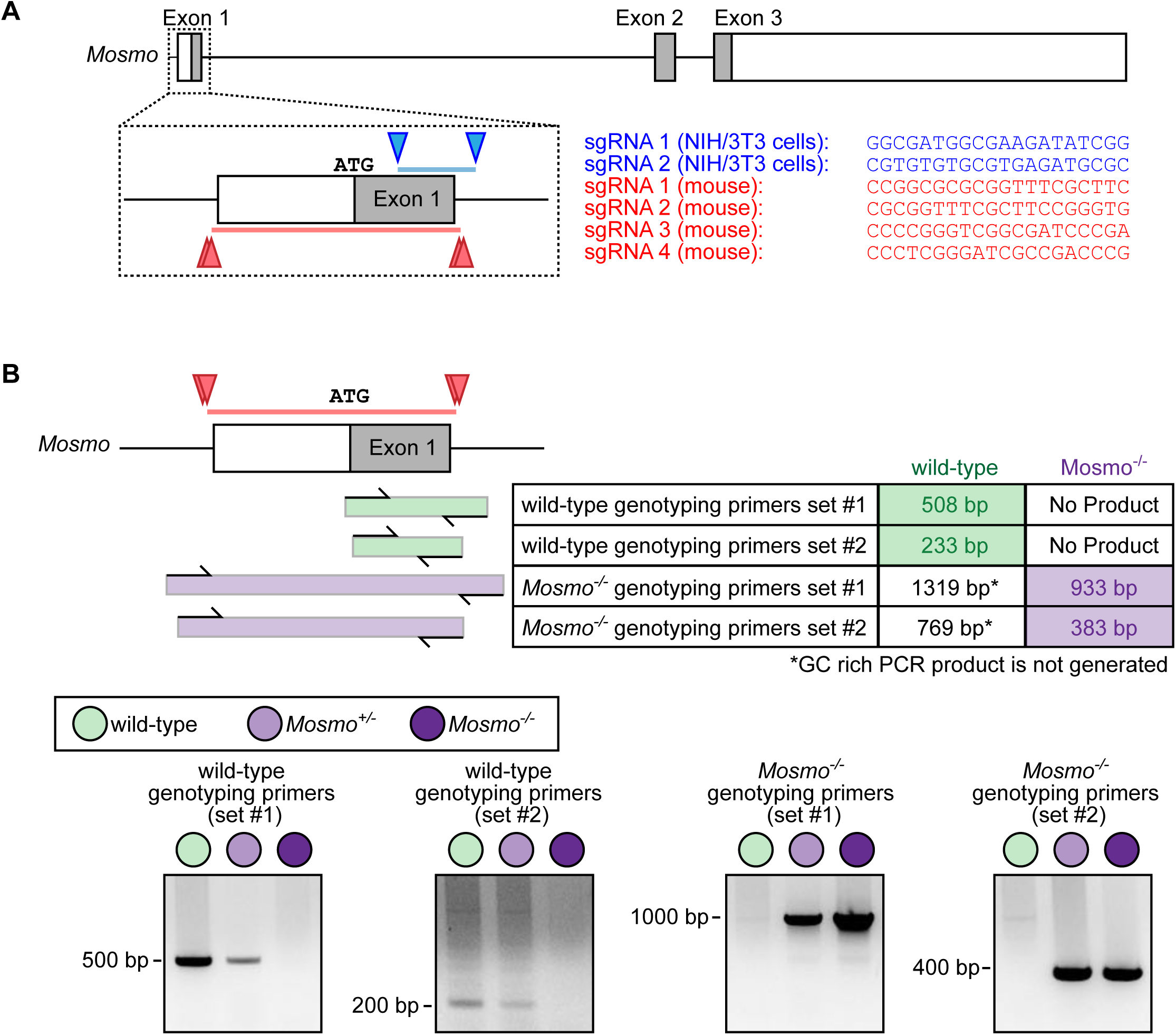
Construction of *Mosmo^-/-^* mice using CRISPR/Cas9 mediated genome editing. **(A)** *Mosmo* knockout (KO) strategy in NIH/3T3 cells and mice. Schematic of the *Mosmo* gene with exons represented as boxes, introns represented as a line, and the coding regions shaded in gray (top). Exon 1 is enlarged (below) with arrows marking the sgRNA guide targets. Guide sequences, targets, and deleted regions in NIH/3T3 cells and mouse embryos are shown in blue and red respectively. **(B)** PCR genotyping strategy to distinguish between wild-type and KO alleles. The *Mosmo^-/-^* mouse has a 386 bp deletion (red line) that includes a removal of the entire first exon (white and gray box). Exon 1 is GC rich and thus a combination of four genotyping primers (located within and outside of the deleted region) were used to determine if the allele has the 386 bp deletion. Representative images of the genotyping PCR are shown below.

**Figure S3:**
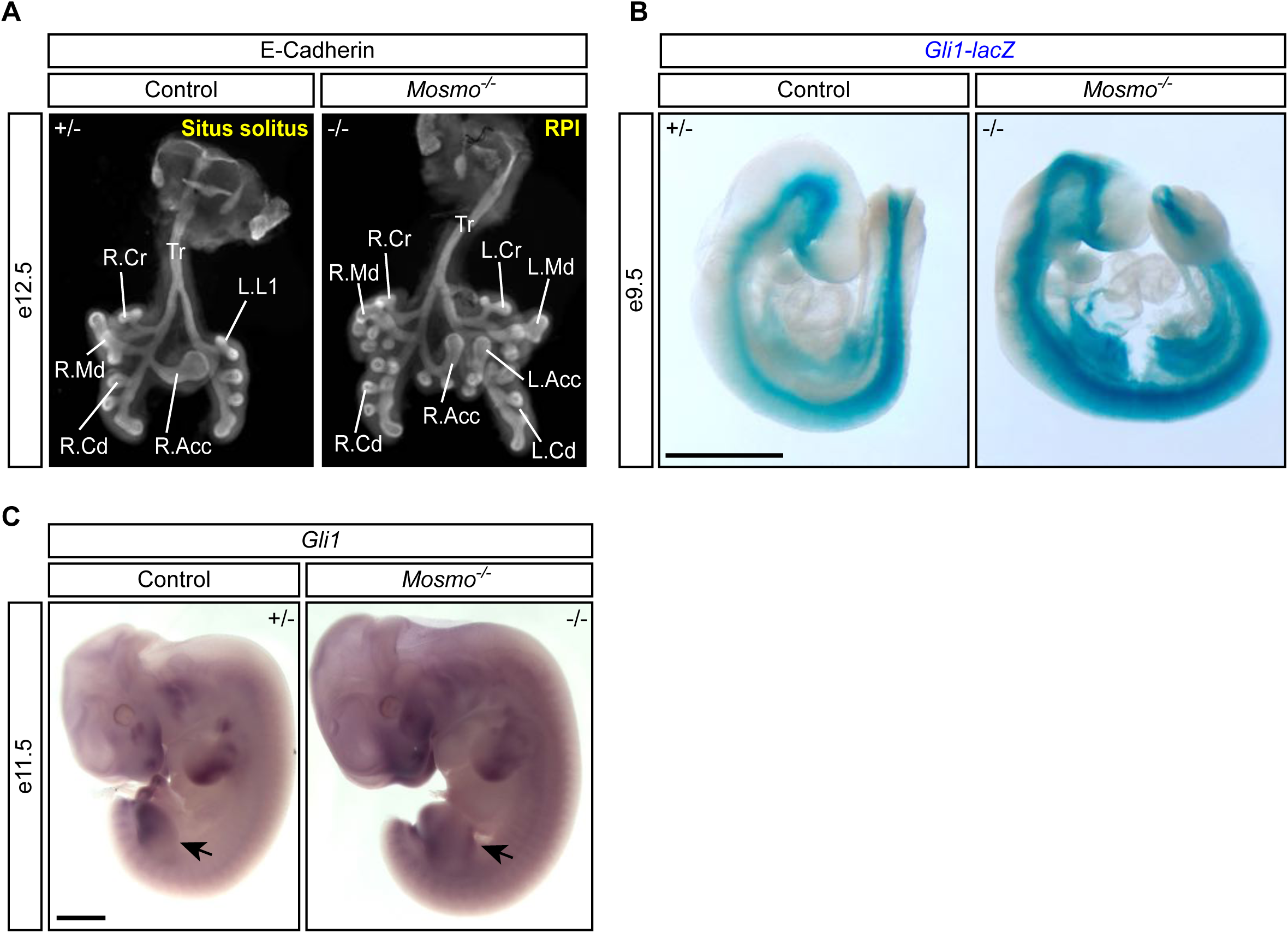
Developmental phenotypes in *Mosmo^-/-^* embryos. **(A)** Whole-mount lungs (ventral view) of e12.5 control (*Mosmo^+/-^*) and *Mosmo*^-/-^ embryos immunostained for E-cadherin to show the airway epithelium. Normal mouse lungs have one lobe on the left (L.L1) and four lobes on the right (R.Acc, right accessory; R.Cr, right cranial; R.Cd, right caudal; R.Md, right middle). *Mosmo*^-/-^ lungs exhibit right pulmonary isomerism (RPI), a duplication of the right lung morphology on the left side (L.Acc, left accessory; L.Cd, left caudal; L.Cr, left cranial; L.Md, left middle). **(B)** Whole-mount e9.5 control (*Mosmo^+/-^*; *Gli1^lacZ/+^*) and knockout (*Mosmo*^-/-^; *Gli1^lacZ/+^*) littermates stained with X-gal to visualize *Gli1-lacZ* expression. Scale bar is 1 mm. **(C)** Whole-mount in situ hybridization was used to assess *Gli1* expression in e11.5 control (*Mosmo^+/-^*) and *Mosmo*^-/-^ littermates. Arrows denote areas of elevated Hh signaling activity in the anterior hindlimb. Scale bar is 1 mm.

**Figure S4:**
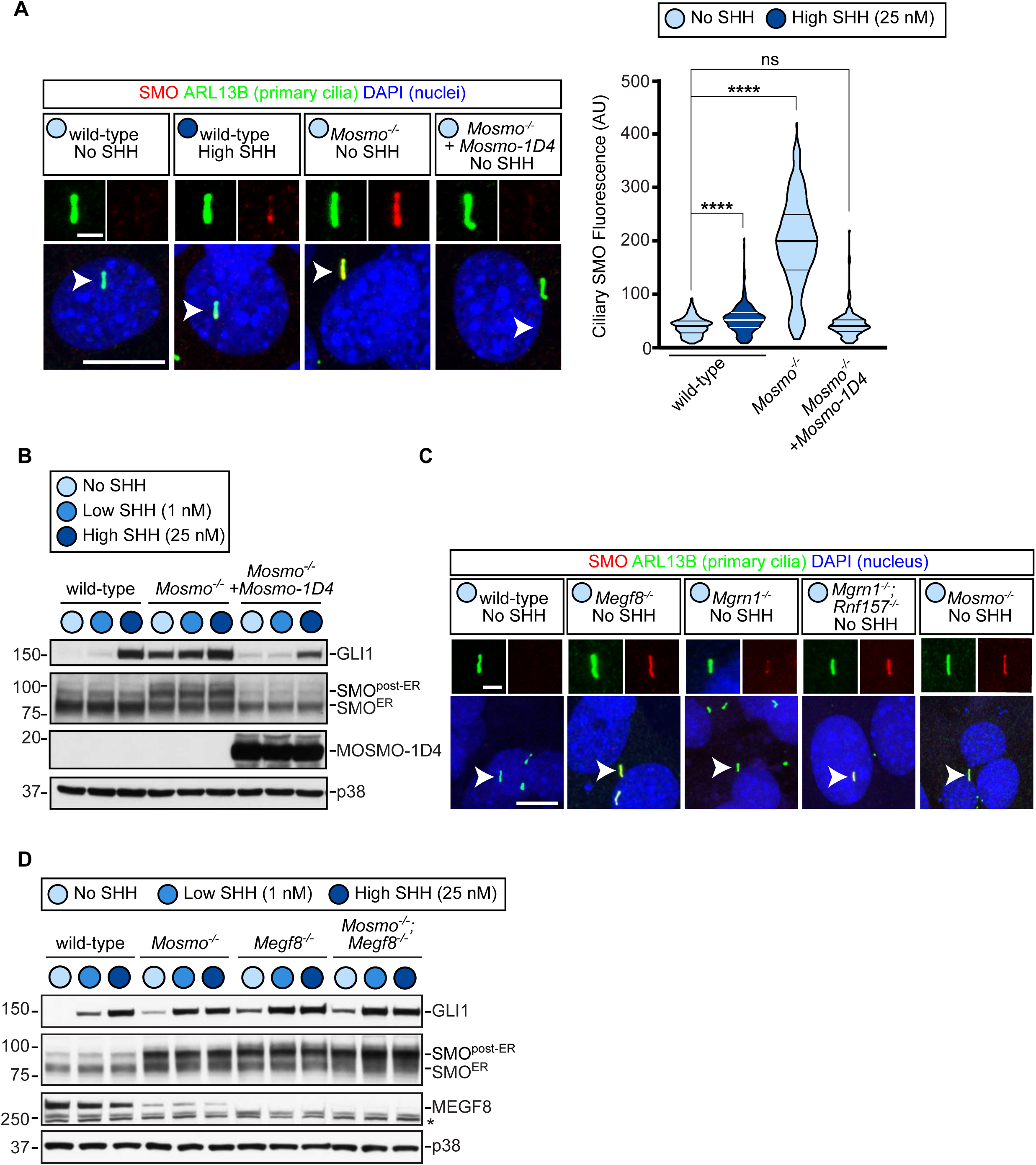
Signaling defects in *Mosmo^-/-^* NIH/3T3 can be corrected by stable re-expression of Mosmo-1D4. **(A-B)** Ciliary SMO abundance in the indicated NIH/3T3 cell lines (wild-type, *Mosmo^-/-^* or *Mosmo^-/-^* with stable re-expression of Mosmo-1D4). Confocal fluorescence microscope images (left) of SMO (red) accumulation at primary cilia (ARL13B, green) in various NIH/3T3 cell lines. Arrowheads identify individual cilia captured in zoomed images above each panel. Nuclei were labeled with DAPI (blue). Ciliary SMO abundance was measured at ∼150 cilia per cell line from multiple such images and used to generate violin plots (right), with horizontal lines denoting the median and interquartile ranges. Statistical significance was determined using the Kruskal-Wallis test; not-significant (ns) > 0.05, ****p value ≤ 0.0001. Scale bars, 10 µm in merged panels and 2 µm in zoomed displays. **(B)** Immunoblots showing GLI1 (as a measure of Hh signaling strength) and SMO abundance in NIH/3T3 cell lines treated with either no, low (1 nM), or high (25 nM) concentrations of SHH. Two populations of SMO are labeled, one localized in the endoplasmic reticulum (ER) and the other localized in post-ER compartments. p38 was used as a loading control. **(C)** Confocal fluorescence microscopy images of SMO (red) localized to the primary cilium (ARL13B, green) in the indicated NIH/3T3 cell lines. Nuclei were labeled with DAPI (blue). Scale bars, 10 µm in merged panels and 2 µm in zoomed displays. **(D)** Immunoblots of GLI1, SMO, and MEGF8 in the indicated NIH/3T3 cell lines treated with varying concentrations of SHH. p38 was used as a loading control. *indicates non-specific bands.

**Figure S5:**
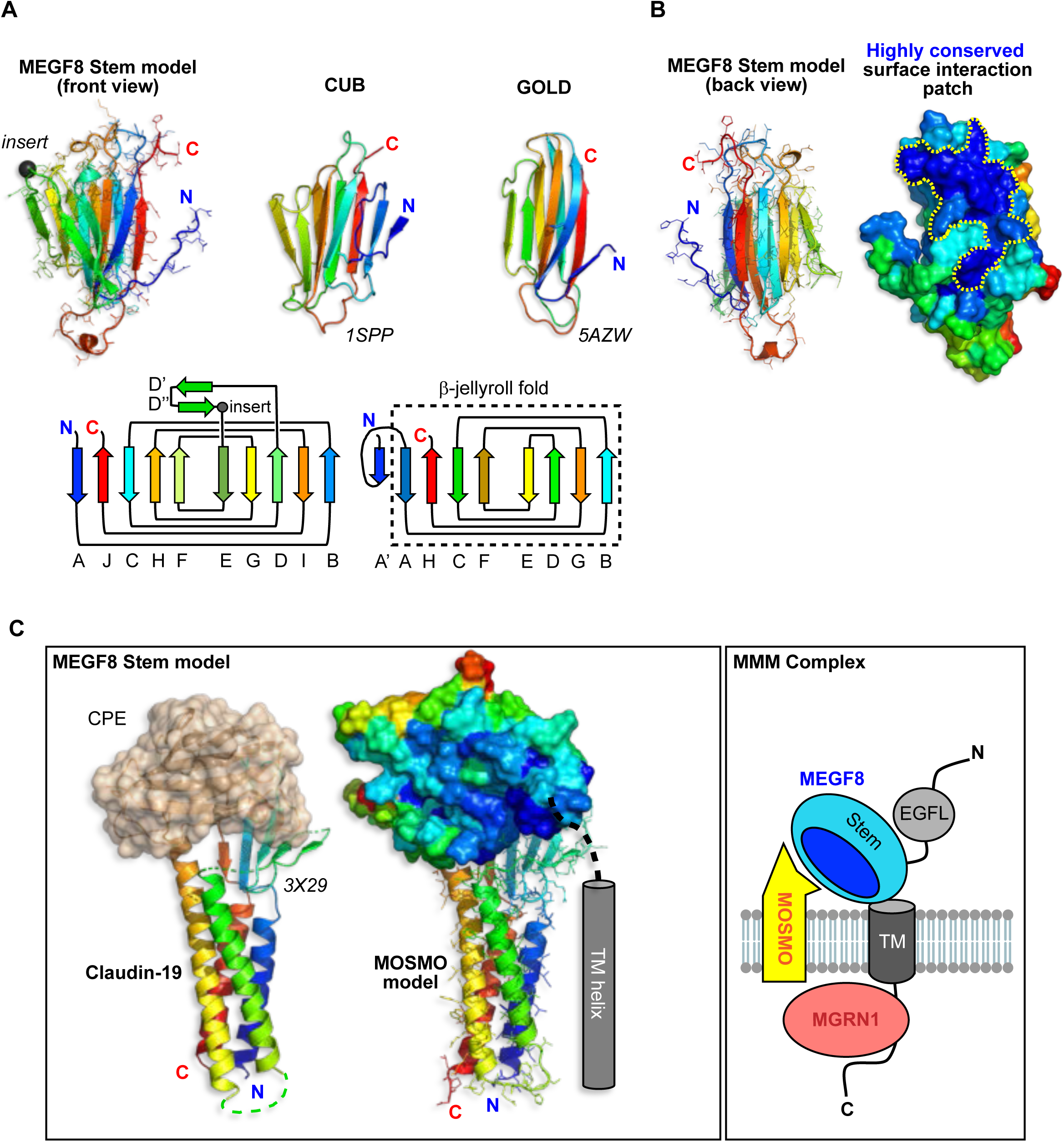
A model of the interaction between the Stem domain of MEGF8 and MOSMO. **(A)** Molecular model of the MEGF8 Stem domain (see Figure 3F) predicted by trRosetta is shown in cartoon form with side chains, the chain color ramped from blue (N-terminus) to red (C-terminus). An unstructured 32 amino acid disordered segment (labeled “insert”) was replaced with a glycine residue (black C sphere) before modelling. Shown to its right are PDBeFOLD superposed CUB and GOLD domains (respectively from PDB files 1SPP and 5AZW) and below are ‘open book’ topology depictions of MEGF8 Stem (left) and the CUB domain (right, with the dotted line showing the β-jellyroll fold overlap with the GOLD domain) with β-strands labeled below. In the Stem domain, we highlight the D’-D” β-hairpin that sprouts between β-strands D and E, and the position of the 32 amino acid unstructured insert (black ball) is noted in the D”-E loop. **(B)** If we rotate the MEGF8 Stem domain model by ∼180°, revealing the ‘back’ β-sheet face (that is free of the D’-D” β-hairpin overhang) and show the surface conservation profile calculated by ConSurf, a highly conserved interaction epitope in dark blue is visible. **(C)** A molecular model of the MOSMO fold was predicted by trRosetta using Claudin structure templates drawn from the PDB and integrating this information with the deep-learning-derived distance and orientation restraints to calculate the final model. To the left of the Claudin-based model for MOSMO is the structure of Claudin-19 docked to the Clostridium perfringens enterotoxin C-terminal domain (CPE, with a β-jellyroll fold related to CUB) from PDB file 3X29. This CPE-Claudin-19 complex offers a template for the interaction of MEGF8’s Stem domain with MOSMO that utilizes the conserved surface patch (colored dark blue in B) in the β-sheet formed by the J, C, H and F strands (see A) of the Stem CUB-like fold. **(D)** A schematic of the MMM complex. The MEGF8 Stem-MOSMO and MEGF8-MGRN1 interactions have been experimentally validated (Figures 3E**-3G** and (Kong et al., 2020)).

**Figure S6:**
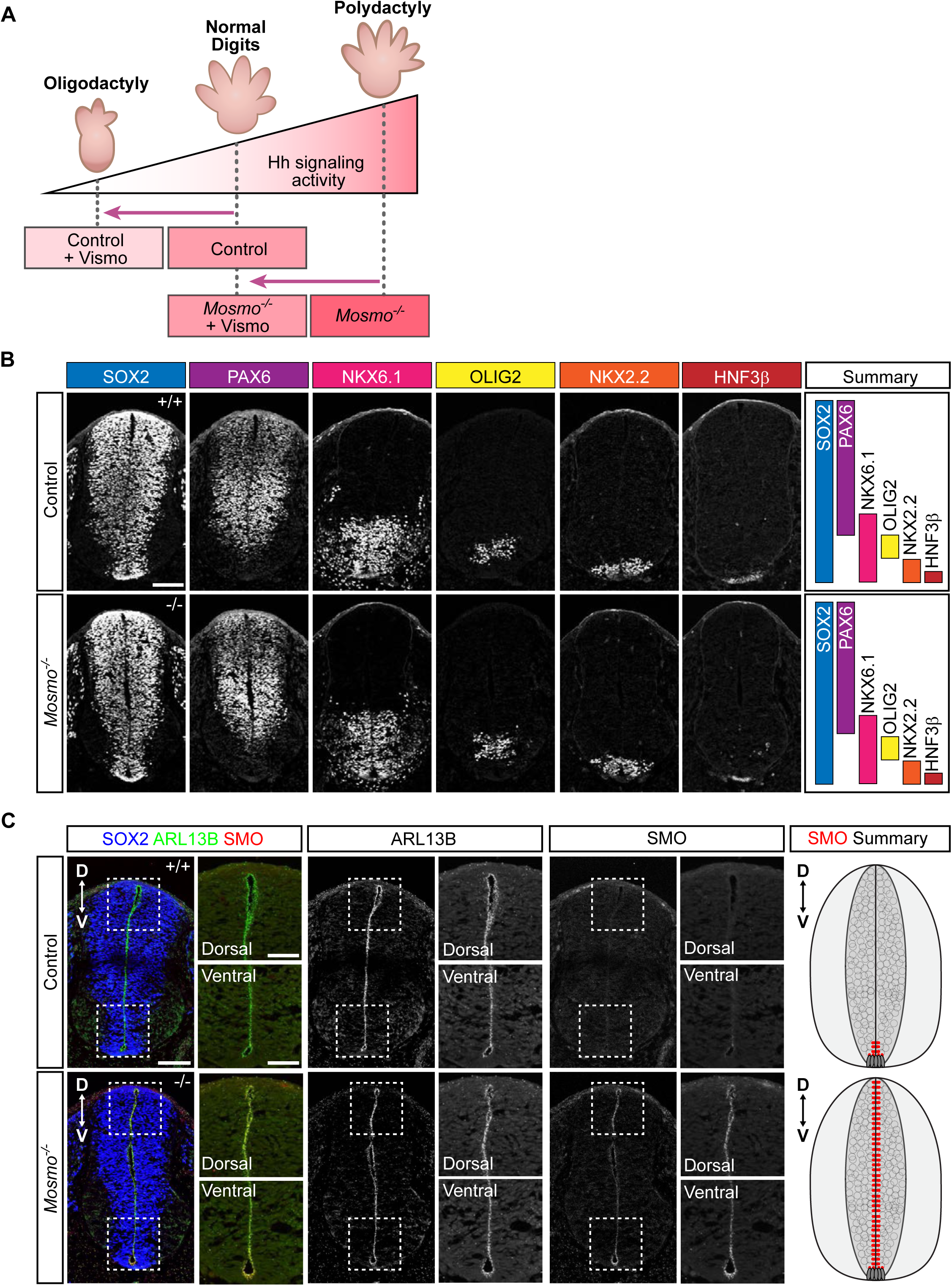
Analysis of spinal cord patterning in control and *Mosmo^-/-^* embryos. **(A)** A proposed model for how a combination of genotype and SMO inhibition by vismodegib influences Hh signaling strength and consequently digit number in developing embryos. **(B-C)** Representative images of transverse sections of e10.5 control (*Mosmo^+/+^* and *Mosmo^+/-^*) and *Mosmo^-/-^* spinal cords. **(B)** Distribution of transcription factors within the developing spinal cord. Scale bar is 100 µm. **(C)** Distribution of SMO (red) within the primary cilia (ARL13B, green) of spinal neural progenitors (SOX2, blue). Scale bars, 100 µm in merged panels and 50 µm in zoomed displays. Quantification available in Figure 6C.

**Table S1.** Related to Figure 1: Genotypes of offspring derived from *Mosmo^+/-^* intercrosses.

| Age | *Mosmo*<sup>+/+</sup> | *Mosmo*<sup>+/-</sup> | *Mosmo*<sup>-/-</sup> | Total | Chi square<br>p-value | Average Litter<br>Size |
| --- | --- | --- | --- | --- | --- | --- |
| *Expected* | 25% | 50% | 25% |  |  |  |
| e9.5-e13.5 | 17 (19.8%) | 43 (50%) | 26 (30.2%) | n = 86 (9 litters) | p = 0.390 | 9.6 |
| e14.5-e18.5 | 24 (32.4%) | 46 (62.2%) | 4 (5.4%) | n = 74 (13 litters) | p = 0.000503 | 5.7 |
| P0-P18 | 12 (28.6%) | 30 (71.4%) | 0 (0%) | n = 42 (8 litters) | p = 0.000685 | 5.3 |

Significant deviation from the expected Mendelian ratio of 1:2:1 was calculated using the chi-square test. Not-significant (ns) > 0.05 and \*\*\*p value ≤ 0.001.

**Table S2:** Related to Figure 1 and 4: Heart, visceral organ situs, and digit number in wild-type (*Mosmo^+/+^*), *Mosmo^-/-^*, and *Mosmo^-/-^*;*Megf8^m/m^* mouse embryos.

| Embryo ID | Body situs | Heart | VA Alignment | Septal Defects | Aortic Arch | Lung Situs | Abdomen Situs | Digitation |
| --- | --- | --- | --- | --- | --- | --- | --- | --- |
| 1 | SS | Lev | CC | None | LAA | 4R:1L | L(St/Sp/P) | Normal |
| 2 | SS | Lev | CC | None | LAA | 4R:1L | L(St/Sp/P) | Normal |
| 3 | SS | Lev | CC | None | LAA | 4R:1L | L(St/Sp/P) | Normal |

***Mosmo*<sup>-/-</sup> embryos**
| Embryo ID | Body situs | Heart | VA Alignment | Septal Defects | Aortic Arch | Lung Situs | Abdomen Situs | Digitation |
| --- | --- | --- | --- | --- | --- | --- | --- | --- |
| 1 | HTX | Mes | TGA | AVSD | LAA | 3R:3L:1M | L(St/Sp/P) | PDD |
| 2 | HTX | Lev | PTA | AVSD | LAA | 3R:3L:1M | L(St/Sp/P) | PDD |
| 3 | HTX | Dex | TGA | AVSD | RAA | 3R:3L:1M | L(St/Sp/P) | PDD |
| 4 | HTX | Mes | TGA | AVSD | RAA | 3R:3L:1M | L(St/Sp/P) | PDD |
| 5 | HTX | Dex | TGA | AVSD | RAA | 1R:4L | L(St/Sp/P) | PDD |
| 6 | SIT | Dex | TGA | AVSD | RAA | 1R:4L | R(St/Sp/P) | PDD |
| 7 | HTX | Dex | DORV | AVSD | RAA | 3R:3L:1M | R(St/Sp/P) | PDD |
| 8 | HTX | Lev | TGA | AVSD | LAA | 3R:3L:1M | L(St/Sp/P) | PDD |
| 9 | HTX | Dex | TGA | AVSD | LAA | 3R:3L:1M | L(St/Sp/P) | PDD |
| 10 | SIT | Dex | TGA | AVSD | RAA | 1R:4L | R(St/Sp/P) | PDD |
| 11 | HTX | Lev | PTA | AVSD | LAA | 3R:3L:1M | L(St/Sp/P) | PDD |

***Mosmo*<sup>-/-</sup>;*Megf8*<sup>m/m</sup> double mutant embryos**
| Embryo ID | Body situs | Heart | VA Alignment | Septal Defects | Aortic Arch | Lung Situs | Abdomen Situs | Digitation |
| --- | --- | --- | --- | --- | --- | --- | --- | --- |
| 1 | HTX | Dex | PTA | - | RAA | 3R:3L:1M | R(St/Sp/P) | PDD |
| 2 | HTX | Dex | TGA | AVSD | RAA | 3R:3L:1M | R(St/Sp/P) | PDD |
| 3 | HTX | Dex | TGA | AVSD | RAA | 3R:3L:1M | R(St/Sp/P) | PDD |
| 4 | HTX | Dex | DORV | VSD | LAA | 3R:3L:1M | L(St/Sp/P) | PDD |
| 5 | HTX | Dex | TGA | AVSD | RAA | 3R:3L:1M | R(St/Sp/P) | PDD |

AVSD, atrioventricular septal defect; CC, concordant ventriculoarterial alignment; Dex, dextrocardia; DORV, double outlet right ventricle; HAA, hypoplastic arch; HTA, hypoplastic transverse heart; HTX, heterotaxy; IAA, interrupted aortic arch; LAA, left aortic arch; Lev, levocardia; L(St/Sp/P), Left sided stomach, spleen, pancreas; Mes, mesocardia; N/A: no digit data collected; NML, normal digit pattern; PDD, preaxial digit duplication; PTA, Persistent truncus arteriosus; RAA, right aortic arch; RPI, right pulmonary isomerism; R(St/Sp/P), Right sided stomach, spleen, pancreas; SIT, situs inversus; SS, situs solitus; SymLiv, symmetric liver; TGA, transposition of the great arteries; VA, ventriculoarterial; VSD, ventricular septal defect; VSD(Ao), ventricular septal defect located below the aorta; 1L, one lung lobe on left side; 4R, 4 lung lobes on right side.

**Table S3:** Related to Figure 1: Lung branching analysis in e12.5 mouse embryos

| Genotype | Total embryos analyzed | Situs solitus | Right pulmonary isomerism (RPI) |  |  |
| --- | --- | --- | --- | --- | --- |
|  |  |  | Complete RPI | Partial RPI |  |
|  |  |  |  | Extra “right” branches on the left | Situs inversus |
| *Mosmo*<sup>+/+</sup> | n = 2 | n = 2/2 (100%) | n = 0/2 (0%) | n = 0/2 (0%) | n = 0/2 (0%) |
| *Mosmo*<sup>+/-</sup> | n = 14 | n = 14/14 (100%) | n = 0/14 (0%) | n = 0/14 (0%) | n = 0/14 (0%) |
| *Mosmo*<sup>-/-</sup> | n = 5 | n = 0/5 (0%) | n = 3/5 (60%) | n = 1/5 (20%) | n = 1/5 (20%) |

**Table S4:** Related to Figure 5: Heart, visceral organ situs, and digit number in control (*Mosmo^+/+^* and *Mosmo^+/-^*) and *Mosmo^-/-^* embryos treated with either vehicle or vismodegib.

***Mosmo*<sup>-/-</sup> embryos with vehicle treatment**
| Embryo ID | Body situs | Heart | VA Alignment | Septal Defects | Aortic Arch | Lung Situs | Abdomen Situs | Digitation |
| --- | --- | --- | --- | --- | --- | --- | --- | --- |
| 1 | HTX | Dex | TGA | AVSD | RAA | 3R:3L:1M | R(St/Sp/P) | PDD |
| 2 | HTX | Lev | TGA | AVSD | LAA | 3R:3L:1M | L(St/Sp/P) | PDD |
| 3 | HTX | Lev | TGA | AVSD | LAA | 3R:3L:1M | L(St/Sp/P) | PDD |
| 4 | HTX | Dex | TGA | AVSD | RAA | 3R:3L:1M | R(St/Sp/P) | N/A |
| 5 | HTX | Dex | TGA | AVSD | RAA | 3R:3L:1M | R(St/Sp/P) | N/A |
| 6 | HTX | Lev | PTA | AVSD | LAA | 3R:3L:1M | L(St/Sp/P) | N/A |

***Mosmo*<sup>+/+</sup> and *Mosmo*<sup>+/-</sup> embryos with 3 Day and 4 Day vismodegib treatment**
| Embryo ID | Body situs | Heart | VA Alignment | Septal Defects | Aortic Arch | Lung Situs | Abdomen Situs | Digitation |
| --- | --- | --- | --- | --- | --- | --- | --- | --- |
| 1 | SS | Lev | PTA | AVSD | LAA | N/A | L(St/Sp/P) | Oligo |
| 2 | SS | Lev | PTA | AVSD | RAA | N/A | L(St/Sp/P) | Oligo |
| 3 | SS | Lev | PTA | AVSD | LAA | N/A | L(St/Sp/P) | Oligo |
| 4 | SS | Lev | PTA | AVSD | LAA | N/A | L(St/Sp/P) | Oligo |
| 5 | SS | Lev | PTA | AVSD | LAA | N/A | L(St/Sp/P) | Oligo |

***Mosmo*<sup>-/-</sup> embryos with 3 Day and 4 Day vismodegib treatment**
| Embryo ID | Vismo (Days) | Body situs | Heart | VA Alignment | Septal Defects | Aortic Arch | Lung Situs | Abdomen Situs | Digitation |
| --- | --- | --- | --- | --- | --- | --- | --- | --- | --- |
| 1 | 3 | HTX | Dex | DORV | AVSD | RAA | 3R:3L:1M | L(St/Sp/P) | Normal/Oligo/PDD |
| 2 | 3 | HTX | Lev | DORV | AVSD | LAA | 3R:3L:1M | L(St/Sp/P) | PDD |
| 3 | 3 | HTX | Lev | DORV | AVSD | LAA | 3R:3L:1M | R(St/Sp/P) | PDD |
| 4 | 3 | HTX | Lev | TGA | AVSD | LAA | 1R:4L | L(St/Sp/P) | Normal/PDD |
| 5 | 3 | HTX | Lev | TGA | AVSD | LAA | 1R:4L | L(St/Sp/P) | Normal/PDD |
| 6 | 3 | HTX | Lev | TGA | AVSD | LAA | 1R:4L | L(St/Sp/P) | Normal/PDD |
| 7 | 4 | HTX | Lev | DORV | AVSD | LAA | 1R:4L | L(St/Sp/P) | Normal/PDD |
| 8 | 4 | HTX | Lev | DORV | AVSD | RAA | 3R:3L:1M | L(St/Sp/P) | Normal/PDD |

Lung situs could not be determined (N/A) in vismodegib treated control embryos due to cystic and hypoplastic lungs. AVSD, atrioventricular septal defect; CC, concordant ventriculoarterial alignment; Dex, dextrocardia; DORV, double outlet right ventricle; HAA, hypoplastic arch; HTA, hypoplastic transverse heart; HTX, heterotaxy; IAA, interrupted aortic arch; LAA, left aortic arch; Lev, levocardia; L(St/Sp/P), Left sided stomach, spleen, pancreas; Mes, mesocardia; N/A: no digit data collected; NML, normal digit pattern; Oligo, oligodactyly; PDD, preaxial digit duplication; PTA, Persistent truncus arteriosus; RAA, right aortic arch; RPI, right pulmonary isomerism; R(St/Sp/P), Right sided stomach, spleen, pancreas; SIT, situs inversus; SS, situs solitus; SymLiv, symmetric liver; TGA, transposition of the great arteries; VA, ventriculoarterial; VSD, ventricular septal defect; VSD(Ao), ventricular septal defect located below the aorta; 1L, one lung lobe on left side; 4R, 4 lung lobes on right side.

**Table S5:**
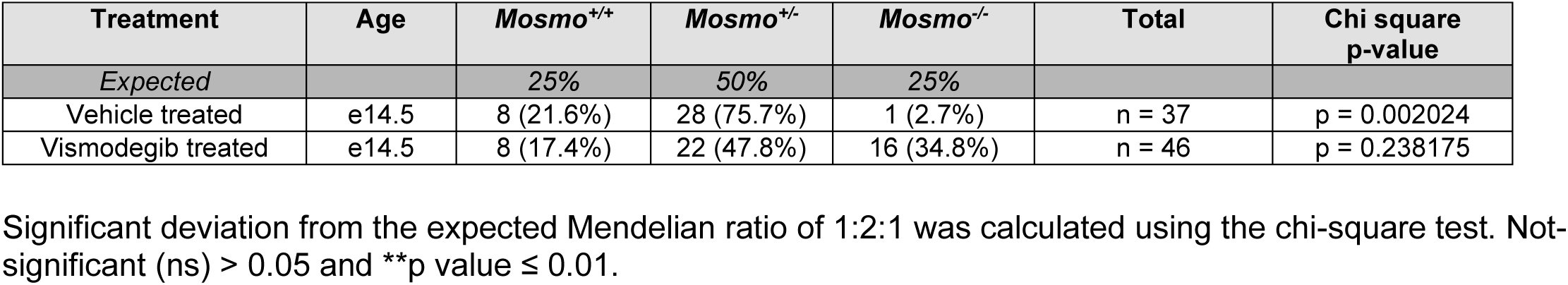
Related to Figure 5: Genotypes of offspring derived from *Mosmo^+/-^* intercrosses treated with vehicle or vismodegib.

